# The transcriptomic signature of obligate parthenogenesis

**DOI:** 10.1101/2021.08.26.457823

**Authors:** Sen Xu, Trung Huynh, Marelize Snyman

## Abstract

Investigating the origin of parthenogenesis through interspecific hybridization can provide insight into how meiosis may be altered by genetic incompatibilities, which is fundamental for our understanding of the formation of reproductive barriers. Yet the genetic mechanisms giving rise to obligate parthenogenesis in eukaryotes remain understudied. In the microcrustacean *Daphnia pulex* species complex, obligately parthenogenetic (OP) isolates emerged as backcrosses of two cyclically parthenogenetic (CP) parental species, *D. pulex* and *D. pulicaria*, two closely related but ecologically distinct species. We examine the genome-wide expression in OP females at the early resting egg production stage, a life-history stage distinguishing OP and CP reproductive strategies, in comparison to CP females of the same stage from the two parental species. Our analyses of the expression data reveal that misregulated genes (underdominant and overdominant genes) are abundant in OP isolates, suggesting widespread regulatory incompatibilities between the parental species. More importantly, underdominant genes (i.e., genes with expression lower than both parentals) in the OP isolates are enriched in meiosis and cell-cycle pathways, indicating an important role of underdominance in the origin of obligate parthenogenesis. Furthermore, metabolic and biosynthesis pathways enriched with overdominant genes (i.e., expression higher than both parentals) are another genomic signature of OP isolates.

## Introduction

Understanding the origin and evolutionary consequences of obligately asexuality (i.e., parthenogenesis) in eukaryotes is critical for deciphering why these lineages are evolutionarily short-lived (Maynard Smith 1978; Bell 1982; Otto 2009). It is well known that the evolutionary consequences of obligate asexuality mainly result from the evolutionarily negligible amount of meiotic recombination in asexual genomes. These include the inability to rapidly adapt due to reduced efficiency of selection (Colegrave 2002; Colegrave et al. 2002; Kaltz and Bell 2002; Poon and Chao 2004; Goddard et al. 2005; Cooper 2007; Kosheleva and Desai 2018), the irreversible accumulation of slightly deleterious mutations (i.e., Muller’s ratcher, Muller 1964; Felsenstein 1974), and the mutational meltdown of asexual populations/lineages (Gabriel et al. 1993; Lynch et al. 1993).

However, the genetic mechanisms underlying the origin of parthenogenetic species remain understudied, despite its potential for informing us about the evolutionary dynamics of asexual taxa. In eukaryotes, obligate parthenogens originate from sexually reproducing ancestors through means of spontaneous mutations, hybridization between closely related sexual lineages, and parasitic infections (Simon et al. 2003). During this sexual-asexual transition, meiosis is altered (e.g., becoming a mitosis-like cell division) to produce chromosomally unreduced gametes. The cytological modifications of meiosis in parthenogens are well studied and differ dramatically between lineages (Stenberg and Saura 2009; Neiman et al. 2014). For example, some lineages engage in apomixis in which the reductional phase of meiosis is skipped to produce unreduced eggs, while others undergo automixis in which the fusion of meiotic products restores the parental ploidy level (Stenberg and Saura 2009). However, due to a lack of understanding of the genetic and molecular mechanisms underlying germline cell division in parthenogens, it remains unknown how these cytological modifications arise.

We suggest that the efforts in resolving this knowledge gap, although still rare, will have implications beyond understanding the evolution of asexuality. For example, understanding how asexuality can originate through hybridization can provide us insights into how meiosis can be disrupted by inter-specific genetic incompatibilities, which is essential to our understanding of how reproductive barriers may arise during speciation. In speciation research, the Dobzhansky-Muller model of hybrid incompatibility posits that genetic elements in two diverging lineages, when placed in the same genetic background, may negatively interact and result in fitness reduction and sterility in hybrids (Dobzhansky 1937; Muller 1940). Substantial empirical evidence supporting Dobzhansky-Muller incompatibilities has emerged over the years (Rawson and Burton 2002; Barbash et al. 2003; Presgraves 2003; Mack and Nachman 2016). However, obligate parthenogens with hybrid origin have not been explicitly considered under this theoretical framework (but see Janko et al. 2018).

Compared to other modes of origin, hybridization is attributed for the largest number of obligate parthenogens by far, with nearly all vertebrate parthenogens having hybrid ancestry (Avise 2015). It is still controversial whether hybridization directly leads to parthenogenesis in the F_1_ generation (Kearney et al. 2009), as only a few attempts succeeded in generating parthenogenetic F_1_s by crossing identified parental lineages found in nature (e.g., Schultz 1973; White et al. 1977; Hotz et al. 1985; Janko et al. 2018). Also, some obligate parthenogens (e.g., *Daphnia*) are backcrosses derived from complex introgression events (Xu et al. 2013; Xu et al. 2015). Because parthenogenetic hybrids are largely incapable of backcrossing with parental lineages, we argue that obligate parthenogenesis in hybrids is analogous to hybrid sterility in reducing gene flow between diverging lineages. Therefore, identifying the genetic elements involved in the hybrid origin of obligate parthenogenesis would greatly expand our understanding of the genetics of hybrid incompatibility.

In this work, we examine whether gene expression plays a role in the hybrid origin of obligate parthenogenesis in the freshwater microcrustacean *Daphnia pulex* species complex. It has long been recognized that differences in gene expression are an important source of phenotypic changes (Wray 2007; Stern and Orgogozo 2008). Gene expression has been examined in various asexual taxa to probe the possible causes of parthenogenesis (Gallot et al. 2012; Hanson et al. 2013; Srinivasan et al. 2014). Nonetheless, for hybrid obligate parthenogens it remains unclear how their gene expression varies relative to the parental species, whether any significant expression changes affect germline cell division, and how regulatory divergences contribute to expression changes.

We address these issues using obligately parthenogenetic (OP) *Daphnia* that are backcrosses of two parental species, the cyclically parthenogenetic (CP) *D. pulex* and *D. pulicaria*, members of the *D. pulex* species complex. *Daphnia* typically reproduces by cyclical parthenogenesis (**Figure 1A**). Under favorable environmental conditions females produce directly developing embryos through parthenogenesis, generating genetically identical daughters (barring *de novo* mutations). However, in unfavorable conditions (e.g., food shortage), some asexual broods become males through environmental sex determination, and females switch to producing haploid eggs. Sexual reproduction between the two produces diapausing, fertilized embryos deposited in a protective case (i.e., ephippium). Interestingly, many populations in the northeast of North America reproduce by obligate parthenogenesis (**Figure 1B**), i.e., reproducing by parthenogenesis in favorable conditions, but also producing ephippial resting embryos by parthenogenesis in deteriorating conditions (Hebert et al. 1993).

**Figure 1.**
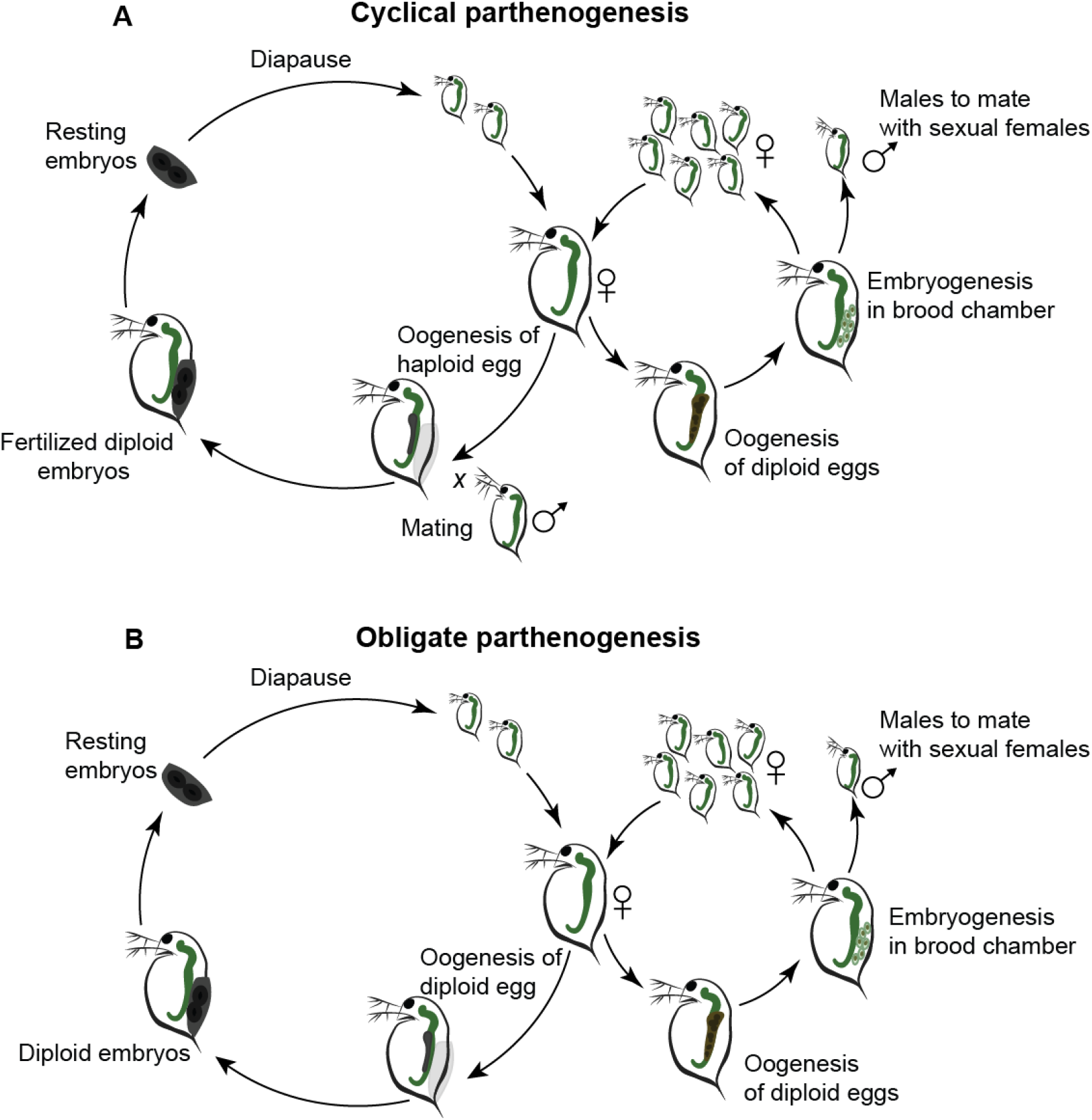
Life history of cyclical parthenogenesis (A) and obligate parthenogenesis (B) in *Daphnia*.

It should be noted that the parthenogenesis of both directly developing embryos and resting embryos in *Daphnia* are most likely achieved by modifications at anaphase I in which no segregation of homologous chromosomes occurs and anaphase I is not followed by formation of daughter cells, i.e., no cytokinesis (Zaffagnini and Sabelli 1972; Hiruta et al. 2010). Therefore, we hypothesize that in hybrid OP *Daphnia* isolates the anaphase promotion complex and spindle checkpoint assembly, which play critical roles in chromosome segregation and cytokinesis (Cooper and Strich 2011; Gorbsky 2015), are affected by aberrant gene expression due to regulatory incompatibility between the parental species, CP *D. pulex* and *D. pulicaria*.

Previous work has revealed that these OP isolates originated through repeated backcrossing between the parental species (Xu et al. 2013; Xu et al. 2015). Genome-wide association mapping showed that the microsatellite and SNPs alleles associated with OP (concentrated on chromosomes 8 and 9) are only present in *D. pulicaria* (a species exclusively inhabiting permanent lake habitats, see below), suggesting that a historical introgression event led to the origin of OP (Lynch et al. 2008; Xu et al. 2015).

Several lines of evidence strongly suggest ecological divergence and genetic incompatibility between these two parental species. As members of the *D. pulex* species complex, CP *D. pulex* and *D. pulicaria* are estimated to have started divergence from 800,000 – 2,000,000 years ago (Colbourne and Hebert 1996; Omilian and Lynch 2009; Cristescu et al. 2012). These two species are morphologically similar (Brandlova et al. 1972) but occupy distinct, overlapping freshwater habitats in North America. *Daphnia pulex* mostly lives in ephemeral fishless ponds, whereas *D. pulicaria* inhabits stratified permanent lakes. These two species show clear physiological differences (Dudycha and Tessier 1999; Caceres and Tessier 2004a, b; Dudycha 2004), indicating strong local adaptation and divergent selection in their distinct habitats.

Although *D. pulex* and *D. pulicaria* can still generate fertile CP F_1_ offspring in laboratory crossing experiments (Heier and Dudycha 2009), some initial evidence points to the presence of both incomplete prezygotic and postzygotic isolation. The prezygotic barrier lies in the effects of photoperiods in triggering sexual reproduction in these two species (Deng 1997), with *D. pulex* switching to sexual reproduction at long day hours (16 hours/day) and *D. pulicaria* switching to sex at short day hours (10 hours/day). On the other hand, postzygotic barriers likely exist as interspecific crosses have lower survival and hatching success than conspecific crosses (Chin et al. 2019).

Despite these reproductive barriers, ample opportunities exist for interspecific introgression between *D. pulex* and *D. pulicaria* when ecological barriers break down (Millette et al. 2020) and phenotypic changes consequently emerged, strongly indicating the widespread presence of genetic incompatibility between them. In addition to obligate parthenogenesis caused by the introgressed *D. pulicaria* alleles in a CP *D. pulex* genetic background, some CP *D. pulex* isolates lost the capability of producing males because they carry an introgressed *D. pulicaria* haplotype at the sex determination loci (Ye et al. 2019). Most likely, because of the genetic incompatibility between the *D. pulex* genetic background and *D. pulicaria* alleles, the sex determination loci in these individuals do not sufficiently respond to environmental cues for initiating the male developmental program in embryos, thus losing the capability of producing male progenies (Ye et al. 2019).

Building upon this rich biological background of the *Daphnia* system, this study examines the gene expression of females at the early stage of resting embryo production (**Figure 1**) to identify the genetic signatures underlying the CP to OP transition. We specifically address (1) the genome-wide expression changes in hybrid OP *Daphnia* in comparison to their parental species and (2) whether aberrant expression pattern affects meiosis and cell cycle (e.g., anaphase promotion complex) in OP individuals and how that may play a role in the origin of obligate parthenogenesis.

## Materials and Methods

### RNA-seq experiment of females at early resting embryo production

Whole-body tissues of mature females at the early stage of resting egg production were collected for 5 OP isolates (Maine344-1, MC08, K09, DB4-4, Maine348-1), 2 CP *D. pulex* (POVI4, PA42), and 2 *D. pulicaria* (AroMoose, LK16) isolates, respectively. These isolates were collected from different geographic locations (Supplementary Table S1). Each isolate has been maintained in lab conditions as an asexual mass culture, which was initiated from a single asexually reproducing female. As gene expression is greatly affected by environmental conditions and by maternal effects, the experimental animals of each isolate were maintained for two asexual generations under 18 °C, and 12:12 (light:dark hours) photoperiod and were fed with the same amount of green algae *Scenedesmus obliquus* as food.

The asexually produced females of the second generations constitute our focal experimental animals. As these animals became mature, we examined these animals daily to search for females engaging in resting egg production. As female *Daphnia* starts resting egg production through normal meiosis in a CP isolate or parthenogenesis in an OP isolate, its ovary shows a characteristic milky color and smooth texture (Hiruta and Tochinai 2014) in contrast to the bulky and dark ovary in females engaging in the parthenogenetic reproduction of directly developing offspring (**Figure 1**). In 1-2 days, an ephippium will start to form in the back of the female due to carapace modification while the ovary continues to develop and become bigger. For the RNA-seq experiments, we collected animals showing the characteristics of early ovary development without sign of ephippium formation, which is what we defined as early resting egg production. Three biological replicates of females in early resting egg production (20-30 individuals) for each *Daphnia* isolate were collected.

For each replicate, the RNA was extracted using the Zymo Insect RNA kit (Zymo Research). The extracted RNA was prepared for short-read sequencing library construction using the NEB NextUltra RNA-seq kit (New England Biolabs). The constructed libraries were sequenced with 150bp paired-end reads on an Illumina HiSeq 2500 and 6000 sequencing platforms. The raw DNA sequence data for this work has been deposited at NCBI SRA under PRJNA726725.

### Inheritance mode of gene expression in OP hybrids

As the parental gene regulatory elements coexist in the OP hybrids, their interaction determines the expression of genes under control. As we hypothesized that the under-expression of genes involved in meiosis and cell cycle is key to the origin of obligate parthenogenesis, we compared the expression level of each gene in the OP hybrids to that of the two CP genotypes representative of each parental lineage (see below). By doing so, we can classify the gene expression inheritance mode of each gene using the following set of rules (**Figure 2A**).

**Figure 2.**
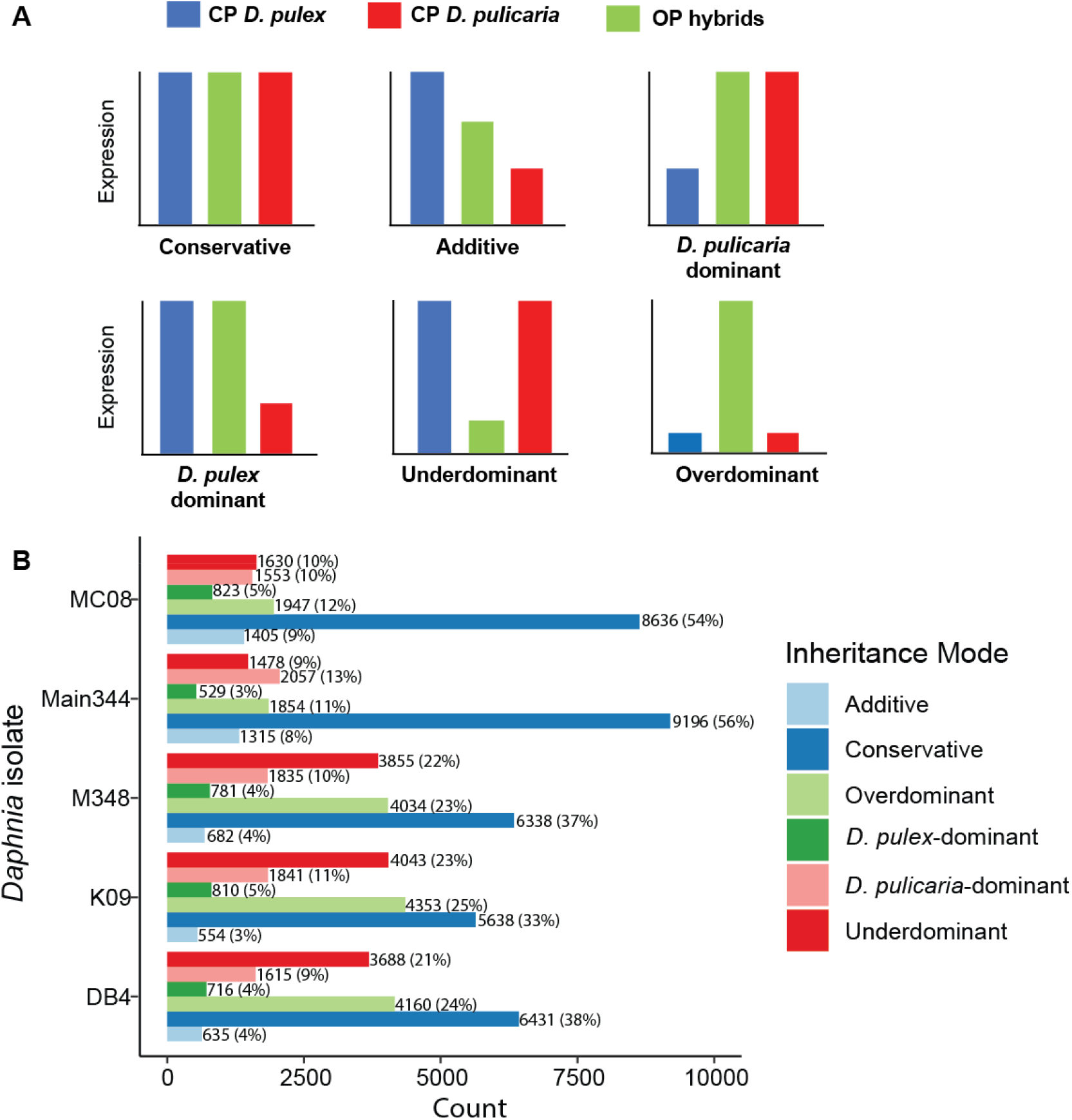
(A) Graphic summary of the criteria for classifying inheritance mode of expression following McManus et al. (2010). (B) Summary of the number of genes in different inheritance mode in five obligately parthenogenetic *Daphnia* isolates. Numbers next to each horizontal bar represent the count of genes, with percentage in the parenthesis.

A gene is considered underdominant if its expression in the OP hybrid is significantly lower than either parental species, whereas a gene is considered overdominant if its expression is higher than either parental species. If the expression of a gene in OP is the same as one parent but significantly different from the other, the inheritance is dominated by one parent, i.e., *D. pulex*- or *D. pulicaria*-dominant. If a gene in OP is expressed at an intermediate level relative to the parentals (i.e., higher than one parental but lower than the other), this gene is considered additive. Lastly, if the expression of a gene in OP is found the same as both parental species, this gene is considered conservative. We consider underdominance and overdominance as misregulated gene expression.

To quantify the gene expression differences between the OP isolates and parental species, the raw reads of these isolates were mapped to the *D. pulex* reference assembly (Ye et al. 2017) using the software STAR (Dobin et al. 2013). As we also planned to do allele-specific analysis (see below) to understand regulatory divergence between parental species, the mapping parameters followed recommendations of Stevenson et al. (2013) for allele-specific RNA-seq analysis using a single reference genome assembly. For all parental species isolates and OP hybrids only the uniquely mapped reads were retained for quantifying gene expression levels. The amount of sequence reads overlapping the gene body of each expressed gene was calculated using the program featureCounts (Liao et al. 2014).

We used the program DESeq2 (Love et al. 2014) to obtain the normalized expression level of each expressed gene in OP hybrids and parental isolates using the median of ratios method. For each gene, we performed pairwise differential expression tests (i.e., Wald test) among one OP hybrid, *D. pulex*, and *D. pulicaria*. Each OP hybrid was individually compared with the parental species, and the parental species were represented by all the biological replicates of the two representative isolates (e.g., *D. pulex* is represented by the biological replicates of POVI4 and PA42). The p-values for differential expression test were corrected for a false discovery rate of 0.05 using the Benjamini-Hochberg method.

A gene’s hybrid expression pattern in an OP hybrid can be delineated if at least two of the three pairwise comparisons are statistically significant. Based on the outcome of these comparisons, we categorized the inheritance mode of genes in each OP hybrid following the rules outlined above (**Figure 2A**). However, for genes with only one statistically significant comparison, we were not able to establish an expression mode and labelled them as ambiguous.

### KEGG (Kyoto Encyclopedia of Genes and Genomes) pathway and GO term analysis

To identify the functional implications of misregulated gene expression (i.e., underdominance and overdominance), we performed a few analyses using annotated KEGG pathways. After mapping *Daphnia* genes into KEGG pathways using the pathway reconstruction tool (https://www.genome.jp/kegg/tool/map_pathway.html), we mapped underdominant and overdominant genes into KEGG pathways in each OP isolate. Within each OP isolate, we calculated a underdominant/overdominant index for each pathway as the ratio of underdominant/overdominant genes. We ranked the underdominant/overdominant index for pathways within each OP isolate and identified the common pathways that were ranked in the top 5 percentile in each OP isolate to understand which pathways were most strongly impacted by misregulation.

We also performed KEGG pathway enrichment analysis and GO (Gene Ontology) term enrichment analysis using the genes that were identified to be underdominant or overdominant in all OP isolates to understand which cellular pathways and functions are disproportionately affected by misregulation. To alleviate the concern that our sampling of the parental species did not cover the entire expression landscape (e.g., some under- and overdominant genes may be false positives), we further selected the underdominant and overdominant genes that were not differentially expressed within either parental species or between the parental species. We identified the differentially expressed genes within the parental *D. pulex* (PA42 *vs* POVI4) and within *D. pulicaria* (LK16 *vs* AroMoose) using DESeq2 (7739 genes in total). Differentially expressed genes (4294 genes) were also identified between the parental *D. pulex* and *D. pulicaria* in DESeq2 using all biological replicates. The final set of underdominant and overdominant genes that went into the KEGG and GO analyses did not vary in expression at within- and between-species level, indicating the importance of their conserved expression level in relation to reproduction in *Daphnia*.

The GO enrichment analysis was performed in the topGO package (Alexa and Rahnenfuhrer 2019), whereas the KEGG pathway enrichment analysis was done using a hypergeometric test with a custom R script.

## Results

### Inheritance mode of gene expression in OP hybrids

We compared the transcript abundance of each expressed gene in OP hybrids to the two parental species to examine its expression inheritance mode. Not surprisingly, conserved expression is the most abundant mode in all OP hybrids, ranging between 33-56% among all isolates (**Figure 2B**). We noted that in the less-well sequenced samples (i.e., MC08 and Main3441), conserved expression is much higher (54% and 56% in MC08 and M3441, respectively) than in the samples of higher sequence coverage (33-38% in M348, K09, DB4), presumably because low sequencing depth reduced our power to detect differential expression in many genes between parental species and OP hybrids. Thus, we suggest that the better-sequenced samples (i.e., M348, K09, DB4) most likely represent an unbiased picture of the inheritance mode of gene expression in OP isolates, whereas in the less-well sequenced samples the proportion of conserved expression is likely inflated.

A striking pattern in the well-sequenced samples (i.e., M348, K09, DB4) is that underdominance and overdominance is the two most abundant expression mode (**Figure 2B**), with overdominance ranging between 23-25% and underdominance between 21-23%. Therefore, misregulated genes (underdominance and overdominance combined) constitute 45-48% of the genes. In the less-well sequenced samples, overdominance ranged between 11-12% and underdominance between 9-10%. However, across the board overdominance is consistently 2-3% higher than underdominance in each OP isolate.

Regarding the other expression modes, additive genes make up 3-9% of the total genes in all the samples. More interestingly, the percentage of *D. pulicaria*-dominant genes (9-13%) is at least two times greater than that of *D. pulex*-dominant genes (3-5%) in each OP hybrid, suggesting more genes resembling the expression pattern in *D. pulicaria* than in *D. pulex*. This is likely because more regulatory elements of *D. pulicaria* show dominance over regulatory elements of *D. pulex*.

### KEGG pathways of top ranked underdominance (UD) and overdominance (OD) index

For each OP isolate, we calculated the ratio of UD and OD genes (i.e., UD and OD index) for each KEGG pathway. This index indicates a possible functional link between misregulated genes and obligate parthenogenesis in *Daphnia*. Notably, we found that the KEGG pathways with a UD index in the top 5 percentile among all 5 OP isolates (**Figure 3A**) were Cell Cycle and Meiosis, suggesting the under-expression of genes in Cell Cycle and Meiosis pathways are key to the origin of obligate parthenogenesis. Pathways involved in embryo development (i.e., Hedgehog signaling Pathway, P53 signaling pathway) were also among the top ranked pathways with regards to UD index, although they were only shared by 4 of the 5 OP isolates (**Figure 3A**).

**Figure 3.**
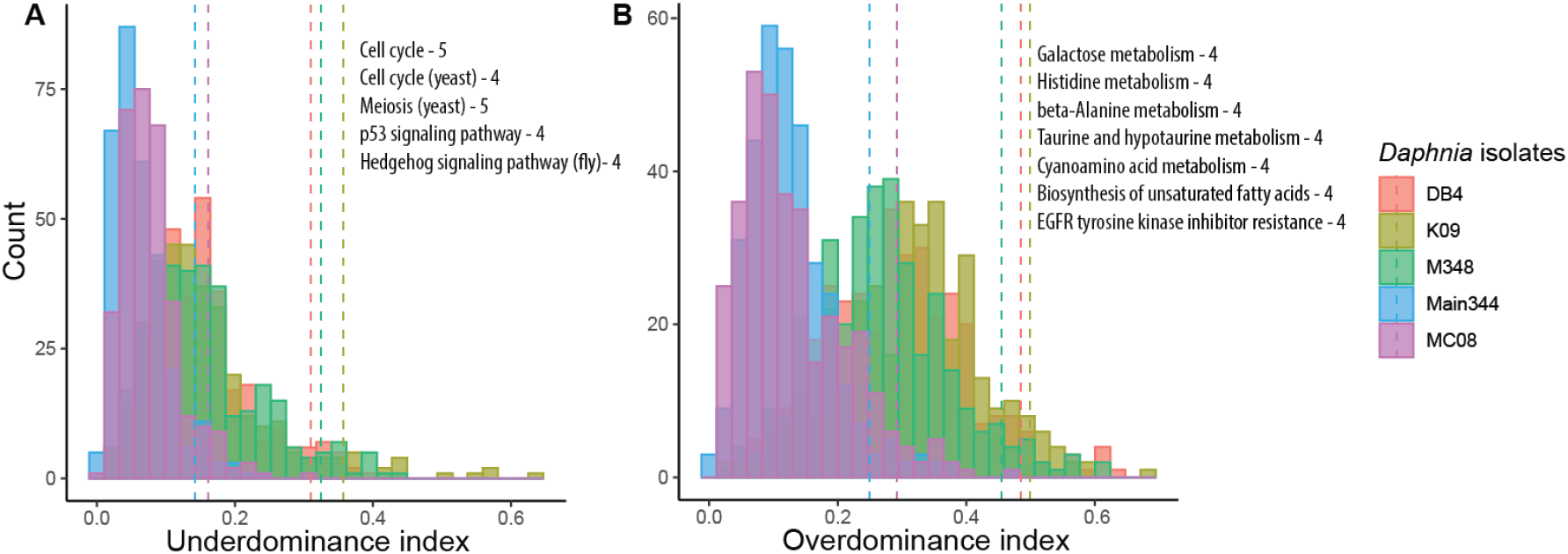
The distribution of underdominance index (A) and overdominance index (B) for KEGG pathways in obligately parthenogenetic *Daphnia* isolates. The vertical dashed lines represent the 95 percentile in each isolate. The KEGG pathways with an index value in the top 5 percentile in at least 4 *Daphnia* isolates (pathway name – number of isolates) are listed.

On the other hand, KEGG pathways of OD index values in the top 5 percentile (**Figure 3B**) were all associated with biosynthesis and metabolism (e.g., Galactose metabolism, Histidine metabolism, Biosynthesis of unsaturated amino acids) with the exception of EGFR tyrosine kinase inhibitor resistance, although it should be noted that these pathways were shared by only 4 of the 5 OP isolates.

### KEGG pathway enrichment analysis

Among genes that are UD (531 genes) or OD (569 genes) in all 5 OP isolates (for gene lists see Supplementary Files 1 and 2), we further selected the genes that have no significant expression variation at within and between species level (for gene lists see Supplementary Files 3 and 4) and used these genes (215 and 265 genes for UD and OD genes, respectively) to perform KEGG pathway enrichment analysis. These gene sets have been mapped to KEGG pathways (see Supplementary Files 5 and 6 for interactive html files) using the software GAEV (Huynh and Xu 2018). This analysis revealed that UD genes were enriched in six pathways related to cell cycle meiosis, and oocyte development (hypergeometric test p < 0.05 with false discovery rate of 0.05), which are Cell Cycle, Meiosis-yeast, Cell Cycle-yeast, Oocyte Meiosis, p53 signaling pathway, and progesterone-mediated oocyte maturation (**Figure 4A**). Furthermore, OD genes were found to be enriched in 23 KEGG pathways (hypergeometric test p < 0.05 with false discovery rate of 0.05), most of which concern metabolism and biosynthesis (**Figure 4B**).

**Figure 4.**
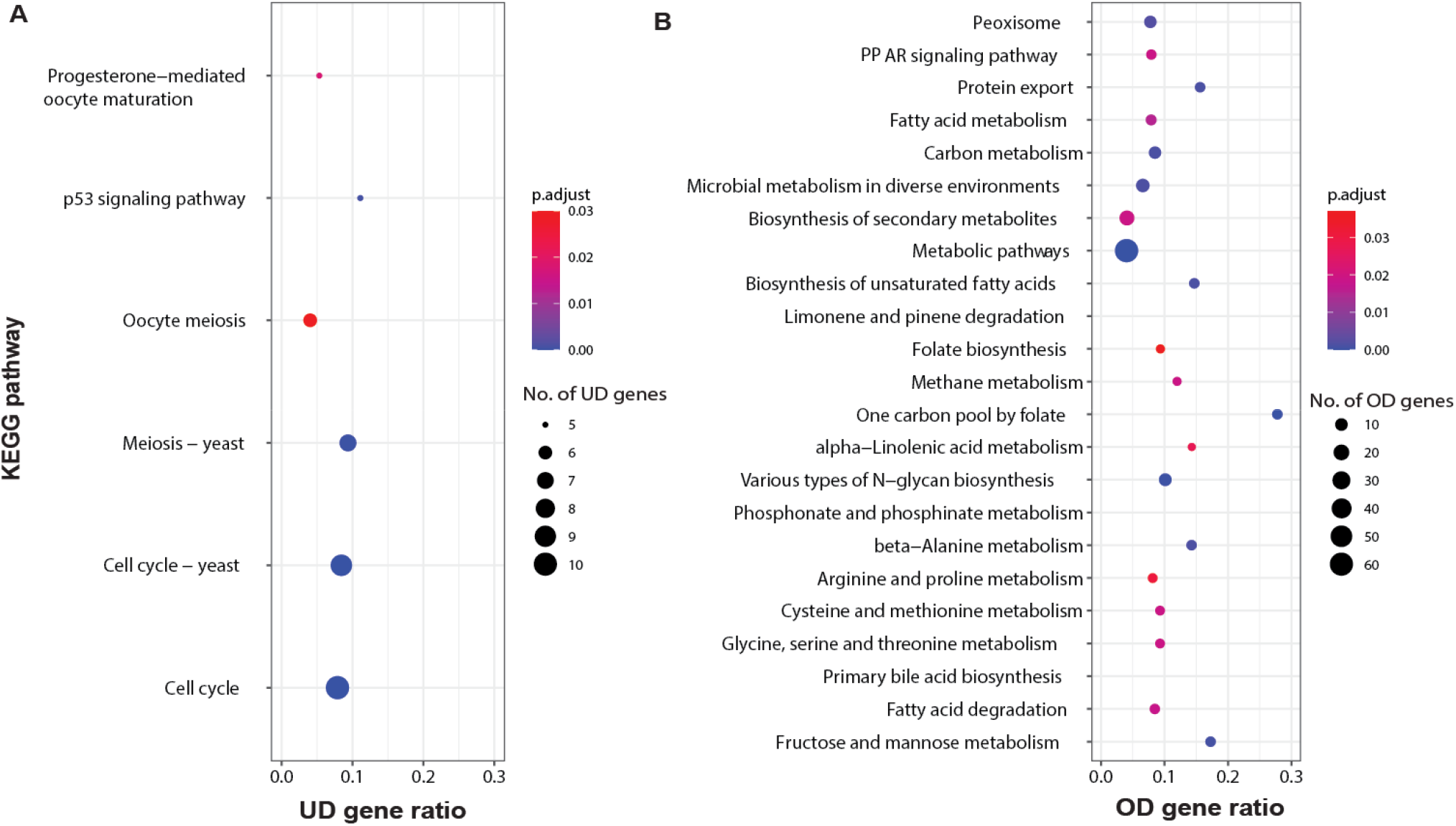
KEGG pathways enriched with underdominant (UD) genes (A) and with overdominant (OD) genes (B). UD/OD gene ratio (horizontal axis) represents the proportion of UD/OD genes in the pathway. p.adjust values are hypergeometric test p values corrected for multiple testing using Benjamini and Hochberg (1995) method.

### GO term enrichment analysis

Nearly all GO terms that were enriched for the UD genes (p < 0.01) were associated with cell cycle, cell division, cell cycle transition, chromosome segregation, and spindle checkpoint assembly (**Supplementary Table S2**). For example, the top five significant GO terms enriched for UD genes were “regulation of mitotic cell cycle”, “cell cycle”, “regulation of cell cycle process”, “mitotic cell cycle process”, and “mitotic cell cycle”. Many GO terms concerning cell cycle transition, chromosome segregation, chromatid separation, spindle assembly checkpoint also showed significant enrichment. On the other hand, the majority of GO terms enriched for OD genes were associated with cell metabolism and biosynthesis (**Supplementary Table S3**), with a notable exception of the top three significant terms concerning protein and macromolecule glycosylation.

### Expression pattern of UD and OD genes

Comparing the transcript abundance of UD and OD genes in each OP isolate to the two parental species, we found that the mean log2-fold expression change of UD genes across five OP isolates is -2.29, whereas the mean log2-fold over-expression of OD genes is 1.93 (**Figure 5A**). The absolute change of UD gene is significantly larger than that of OD genes (two sample t-test, p < 2 × 10^−16^).

**Figure 5.**
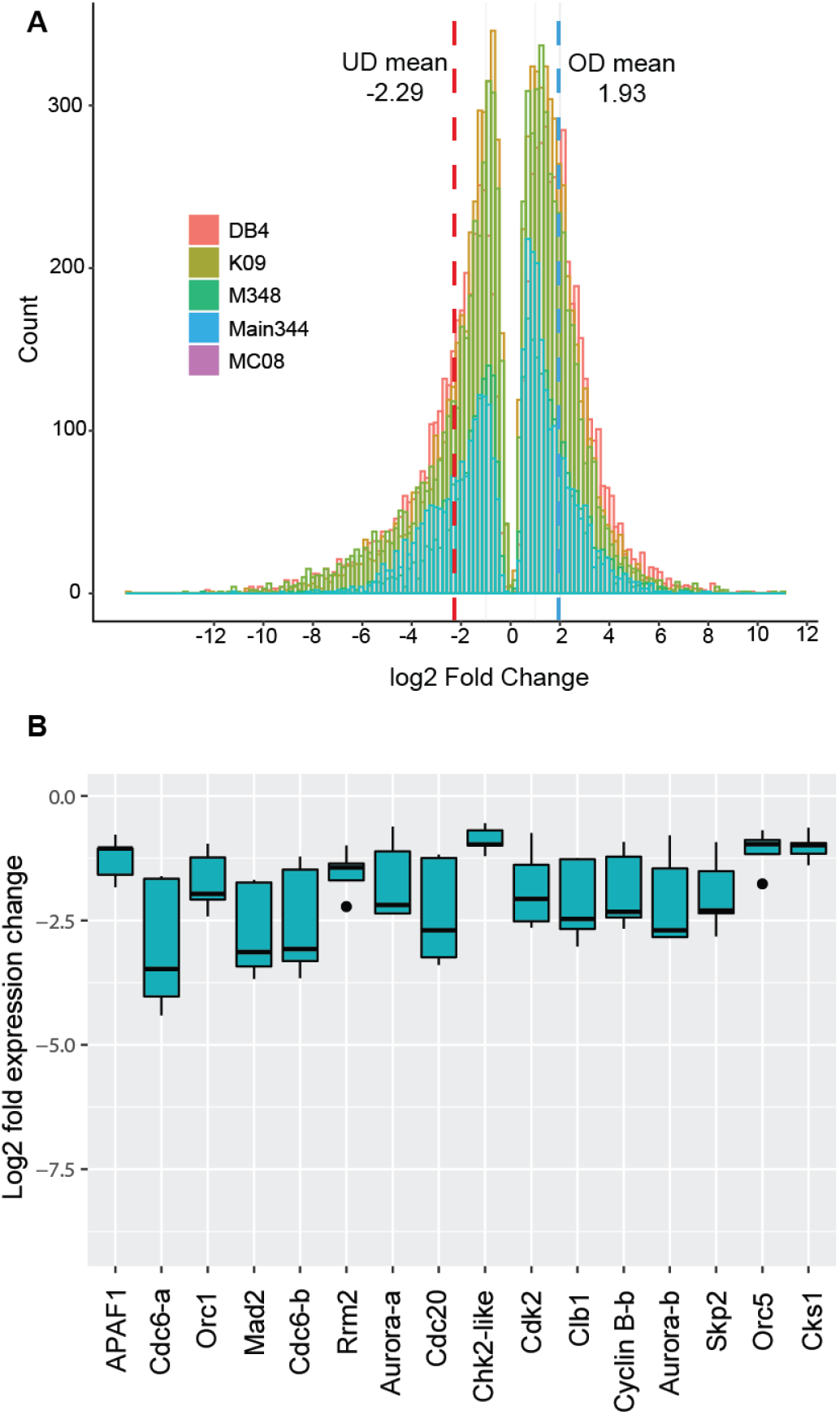
(A) Histograms of log2-fold change of underdominant (UD) and overdominant (OD) genes in obligately parthenogenetic *Daphnia* isolates, with mean value indicated by dashed lines. (B) Log2-fold expression change of meiosis and cell-cycle genes that were underdominant in all examined obligately parthenogenetic *Daphnia* isolates.

As the misregulated genes in meiosis and cell cycle pathways were predominantly UD, We identified several UD genes that are shared by all OP isolates and do now show expression variation at both within and between species. Their log2-fold under-expression relative to the parental species ranged between -0.5 and -4.5 (**Figure 5B**) in OP isolates. Many of these genes are key players in the anaphase promotion complex (e.g., Cdc20), spindle checkpoint assembly (e.g., Mad2), and cell cycle control (e.g., cell cyclin B).

## Discussion

Examining gene expression in obligate parthenogens of hybrid origin in comparison with sexual parental species is an effective approach to understanding how expression changes and interspecific genetic incompatibility may play a role in the origin of asexuality. Indeed, our analyses of the gene expression changes in OP *Daphnia* relative to the CP parental species, *D. pulex* and *D. pulicaria*, revealed several important genetic modifications likely involved in the origin of obligate parthenogenesis.

First, misregulated gene expression (i.e., underdominant and overdominant combined) is abundant in OP *Daphnia* (i.e., up to 45-48%), while conserved expression has the highest number of genes out of any single inheritance mode of expression (**Figure 2B**). Moreover, the number of overdominant genes in OP *Daphnia* is consistently 2-3% higher than that of underdominant genes in all the examined samples. Clearly, compared to either parental species, OP *Daphnia* harbors genes with significantly altered expression patterns. This stands in stark contrast with the observation in the hybrid genome of asexual *Cobitis* loaches, where dominance by one parental species accounts for the expression in 83-89% genes and mis-regulation appears to be rare (Bartos et al. 2019). Therefore, it remains to be seen how the amount of misregulated genes in *Daphnia* compares to other hybrid obligate parthenogens and whether there are consistently more overdominant *vs*. underdominant genes because this kind of data is lacking from other obligate parthenogens.

Second, our analyses demonstrate that the under-expression of meiosis and cell-cycle genes and over-expression of metabolic genes is a key genomic signature of parthenogenesis in *Daphnia*. This is manifested in the significant enrichment of underdominant genes in meiosis and cell-cycle pathways and GO terms and enrichment of overdominant genes in metabolic pathways and GO terms in the early resting egg production stage of OP *Daphnia*. It should be noted that the set of overdominant and underdominant genes used in the KEGG pathway and GO term analyses are very conservative because we kept only genes that did not have variable expression at within and between species levels. Although one may argue that we only sampled two genotypes of each parental species, these overdominant and underdominant genes having conserved expression at within- and between-species level strongly suggests that these genes would most likely remain to be identified as underdominant or overdominant when more parental genotypes are included for analysis.

This transcriptomic signature is not only a characteristic of early resting egg production through parthenogenesis in OP hybrids, but also identified in the females at early stage of asexually producing directly developing embryos in CP *D. pulex* and CP *D. pulicaria* when compared to the early stage of meiotically producing haploid eggs (Huynh et al. unpublished data). We therefore hypothesize that the under-expression of meiosis and cell-cycle genes plays an important role in the origin of obligate parthenogenesis in hybrid OP *Daphnia*. On the other hand, the over-expression of metabolic genes may be a trigger of parthenogenetic reproductive pathways or simply a response to environmental cues. Testing these hypotheses would entail detailed molecular genetic analyses of these genes in relation to obligately parthenogenetic reproduction. We suspect that the causal factors would likely involve only a few key under- and overdominant genes affecting key pathways, whereas many of the other identified genes could be of secondary consequences.

One noteworthy underdominant gene (i.e., Cdc20) has also been identified as candidate genes involved in the origin of obligate parthenogenesis in *Daphnia* (Eads et al. 2012; Xu et al. 2015), whereas another underdominant gene Mad2 is an important member of the spindle checkpoint assembly. As the parthenogenetic production of resting embryos in OP *Daphnia* is most likely achieved through a modified form of meiosis (Zaffagnini and Sabelli 1972) where segregation is suppressed and anaphase I does not lead to cytokinesis, the under-expression of these genes in OP *Daphnia* may very well be essential to these modifications. Cdc20 is a key cell cycle regulator (Yu 2007) and its under-expression may play a more significant role than other genes. Although our results are based on RNA-seq of whole-body tissue, this idea draws support from emerging molecular evidence. For example, the under-expression of Cdc20 in mouse results in failed segregation of chromosomes in oocytes (Jin et al. 2010). This idea is worth of further investigation at the tissue-specific and cellular level.

Third, the genetic mechanisms underlying the misregulation of the identified genes remain to be examined. Identifying the genetic loci and variants regulating the expression of these genes through eQTL (expression quantitative trait loci) mapping would be helpful for us to answer questions such as whether the misregulation is a direct consequence of the interaction between the introgressed *D. pulicaria* alleles with *D. pulex* genetic background.

Lastly, we emphasize that OP in *D. pulex* is most likely a polygenic trait (Xu et al. 2015). There are other genetic components involved in OP that need to be further investigated, e.g., the suppression of recombination during homologous pairing and the bypass of oocyte development arrest without fertilization. Hopefully, future studies can identify the possible candidate genes. Most importantly, we believe that molecular functional characterization of the roles of the identified candidate genes in OP is much needed, which can provide a novel perspective on the origin of parthenogenesis and on the genetic incompatibility in closely related species.

## Acknowledgements

We thank A. Hall for his help with the experiments. This work is supported by NIH MIRA grant R35GM133730 to SX. This research used computational resources supported by the National Science Foundation under Grant Nos. DBI-1062432 2011, ABI-1458641 2015, and ABI-1759906 2018 to Indiana University. Any opinions, findings, and conclusions or recommendations expressed in this material are those of the authors and do not necessarily reflect the views of the National Science Foundation, the National Center for Genome Analysis Support, or Indiana University.

