## Supplementary material for "The transcriptomic signature of obligate parthenogenesis": Supplementary File 5-original_UD(red)_OD(green).html

  

Pathway Sorted by Number of Associated Genes

| Pathway  Pathway |
| --- |
| PI3K-Akt signaling pathway(15, 14.56%) |
| Biosynthesis of cofactors(15, 18.29%) |
| Cell cycle(14, 19.18%) |
| Viral carcinogenesis(11, 13.92%) |
| Lysosome(11, 17.46%) |
| Oocyte meiosis(10, 16.67%) |
| Apoptosis(10, 17.86%) |
| Human T-cell leukemia virus 1 infection(10, 11.11%) |
| Peroxisome(10, 21.74%) |
| Estrogen signaling pathway(9, 18.75%) |
| Proteoglycans in cancer(9, 9.68%) |
| Hepatitis B(9, 15.52%) |
| Transcriptional misregulation in cancer(9, 12.50%) |
| Cell cycle - yeast(9, 14.75%) |
| Focal adhesion(9, 11.54%) |
| Calcium signaling pathway(8, 11.27%) |
| Progesterone-mediated oocyte maturation(8, 17.02%) |
| Prostate cancer(8, 20.51%) |
| MicroRNAs in cancer(8, 13.33%) |
| Protein digestion and absorption(8, 24.24%) |
| Epstein-Barr virus infection(7, 9.86%) |
| Choline metabolism in cancer(7, 18.92%) |
| Carbon metabolism(7, 9.59%) |
| FoxO signaling pathway(7, 12.50%) |
| Meiosis - yeast(7, 14.89%) |
| Hepatocellular carcinoma(7, 10.45%) |
| Relaxin signaling pathway(7, 12.73%) |
| Protein export(7, 33.33%) |
| Glycolysis / Gluconeogenesis(7, 21.88%) |
| Cushing syndrome(6, 8.70%) |
| Thyroid hormone synthesis(6, 18.18%) |
| Ras signaling pathway(6, 7.79%) |
| Pathogenic Escherichia coli infection(6, 9.38%) |
| Measles(6, 13.64%) |
| Influenza A(6, 13.64%) |
| Antigen processing and presentation(6, 37.50%) |
| Ubiquitin mediated proteolysis(6, 7.32%) |
| Vibrio cholerae infection(6, 20.00%) |
| Small cell lung cancer(6, 15.79%) |
| ECM-receptor interaction(6, 27.27%) |
| Hippo signaling pathway - fly(6, 13.64%) |
| Fructose and mannose metabolism(6, 31.58%) |
| Signaling pathways regulating pluripotency of stem cells(6, 11.54%) |
| p53 signaling pathway(6, 20.00%) |
| Porphyrin and chlorophyll metabolism(6, 35.29%) |
| Autophagy - animal(5, 6.25%) |
| Cellular senescence(5, 7.46%) |
| NOD-like receptor signaling pathway(5, 10.00%) |
| Parathyroid hormone synthesis, secretion and action(5, 11.11%) |
| Growth hormone synthesis, secretion and action(5, 10.00%) |
| Pancreatic secretion(5, 12.20%) |
| Spinocerebellar ataxia(5, 6.02%) |
| Human cytomegalovirus infection(5, 6.58%) |
| MAPK signaling pathway(5, 5.56%) |
| Salmonella infection(5, 4.95%) |
| HIF-1 signaling pathway(5, 12.20%) |
| Lysine degradation(5, 13.89%) |
| Tight junction(5, 7.58%) |
| Regulation of actin cytoskeleton(5, 6.85%) |
| AMPK signaling pathway(5, 9.09%) |
| Adipocytokine signaling pathway(5, 20.00%) |
| Fatty acid metabolism(5, 15.15%) |
| Chemokine signaling pathway(5, 8.93%) |
| Wnt signaling pathway(5, 6.94%) |
| Alcoholism(5, 10.64%) |
| One carbon pool by folate(5, 45.45%) |
| EGFR tyrosine kinase inhibitor resistance(5, 17.24%) |
| Rap1 signaling pathway(5, 6.76%) |
| Folate biosynthesis(5, 29.41%) |
| Phagosome(5, 9.80%) |
| Hepatitis C(5, 12.50%) |
| Amino sugar and nucleotide sugar metabolism(5, 16.67%) |
| Legionellosis(5, 21.74%) |
| Apoptosis - fly(5, 12.20%) |
| mTOR signaling pathway(5, 5.75%) |
| Metabolism of xenobiotics by cytochrome P450(5, 55.56%) |
| Drug metabolism - cytochrome P450(5, 55.56%) |
| Chemical carcinogenesis(5, 33.33%) |
| Gastric cancer(5, 9.09%) |
| Glycosphingolipid biosynthesis - lacto and neolacto series(5, 45.45%) |
| Dorso-ventral axis formation(5, 22.73%) |
| Cysteine and methionine metabolism(5, 16.13%) |
| Other types of O-glycan biosynthesis(5, 33.33%) |
| Glycine, serine and threonine metabolism(5, 17.86%) |
| Tyrosine metabolism(5, 27.78%) |
| Starch and sucrose metabolism(5, 23.81%) |
| cGMP-PKG signaling pathway(4, 6.78%) |
| Vascular smooth muscle contraction(4, 9.30%) |
| GnRH signaling pathway(4, 10.00%) |
| Shigellosis(4, 3.88%) |
| Human immunodeficiency virus 1 infection(4, 5.06%) |
| PD-L1 expression and PD-1 checkpoint pathway in cancer(4, 12.90%) |
| Toxoplasmosis(4, 11.11%) |
| Tuberculosis(4, 8.00%) |
| Herpes simplex virus 1 infection(4, 7.27%) |
| Th17 cell differentiation(4, 16.00%) |
| Thyroid hormone signaling pathway(4, 7.02%) |
| Purine metabolism(4, 5.97%) |
| PPAR signaling pathway(4, 14.29%) |
| Non-alcoholic fatty liver disease(4, 4.76%) |
| Necroptosis(4, 11.11%) |
| Fluid shear stress and atherosclerosis(4, 9.09%) |
| Biosynthesis of unsaturated fatty acids(4, 36.36%) |
| mRNA surveillance pathway(4, 7.02%) |
| cAMP signaling pathway(4, 5.88%) |
| MAPK signaling pathway - fly(4, 6.06%) |
| TNF signaling pathway(4, 10.81%) |
| Galactose metabolism(4, 22.22%) |
| ErbB signaling pathway(4, 12.12%) |
| Phospholipase D signaling pathway(4, 7.55%) |
| Th1 and Th2 cell differentiation(4, 17.39%) |
| AGE-RAGE signaling pathway in diabetic complications(4, 11.76%) |
| Glioma(4, 14.81%) |
| Non-small cell lung cancer(4, 16.00%) |
| Neuroactive ligand-receptor interaction(4, 6.67%) |
| Insulin resistance(4, 8.89%) |
| Fatty acid degradation(4, 15.38%) |
| Various types of N-glycan biosynthesis(4, 15.38%) |
| Platinum drug resistance(4, 14.29%) |
| Breast cancer(4, 7.14%) |
| Glycerolipid metabolism(4, 14.81%) |
| Glutathione metabolism(4, 16.67%) |
| Drug metabolism - other enzymes(4, 18.18%) |
| Glycerophospholipid metabolism(4, 10.00%) |
| Tryptophan metabolism(4, 16.00%) |
| Biosynthesis of amino acids(4, 9.52%) |
| Retinol metabolism(4, 36.36%) |
| beta-Alanine metabolism(4, 23.53%) |
| Phosphatidylinositol signaling system(3, 7.14%) |
| Gap junction(3, 8.82%) |
| Retrograde endocannabinoid signaling(3, 4.41%) |
| Cholinergic synapse(3, 7.14%) |
| Dopaminergic synapse(3, 6.67%) |
| Oxytocin signaling pathway(3, 5.26%) |
| Glucagon signaling pathway(3, 6.98%) |
| Kaposi sarcoma-associated herpesvirus infection(3, 5.77%) |
| Notch signaling pathway(3, 12.00%) |
| Inositol phosphate metabolism(3, 8.33%) |
| T cell receptor signaling pathway(3, 8.33%) |
| Neurotrophin signaling pathway(3, 5.88%) |
| Yersinia infection(3, 4.92%) |
| Longevity regulating pathway - worm(3, 7.69%) |
| Axon regeneration(3, 6.25%) |
| Fc gamma R-mediated phagocytosis(3, 8.33%) |
| Plant-pathogen interaction(3, 27.27%) |
| IL-17 signaling pathway(3, 11.11%) |
| Adherens junction(3, 8.33%) |
| JAK-STAT signaling pathway(3, 14.29%) |
| Melanogenesis(3, 7.50%) |
| Aminoacyl-tRNA biosynthesis(3, 11.54%) |
| Osteoclast differentiation(3, 10.71%) |
| Carbohydrate digestion and absorption(3, 21.43%) |
| MAPK signaling pathway - yeast(3, 10.34%) |
| Adrenergic signaling in cardiomyocytes(3, 6.98%) |
| Hypertrophic cardiomyopathy(3, 12.50%) |
| Axon guidance(3, 4.55%) |
| Natural killer cell mediated cytotoxicity(3, 13.64%) |
| Fc epsilon RI signaling pathway(3, 14.29%) |
| Epithelial cell signaling in Helicobacter pylori infection(3, 9.38%) |
| Complement and coagulation cascades(3, 21.43%) |
| GABAergic synapse(3, 9.68%) |
| RIG-I-like receptor signaling pathway(3, 17.65%) |
| Valine, leucine and isoleucine degradation(3, 8.57%) |
| Fat digestion and absorption(3, 23.08%) |
| Hippo signaling pathway(3, 5.26%) |
| Pentose and glucuronate interconversions(3, 21.43%) |
| Insulin signaling pathway(3, 5.17%) |
| Insect hormone biosynthesis(3, 30.00%) |
| Endocrine resistance(3, 6.82%) |
| Amoebiasis(3, 10.00%) |
| Rheumatoid arthritis(3, 14.29%) |
| Colorectal cancer(3, 7.89%) |
| Acute myeloid leukemia(3, 13.64%) |
| Alanine, aspartate and glutamate metabolism(3, 12.50%) |
| Amphetamine addiction(3, 10.71%) |
| Glyoxylate and dicarboxylate metabolism(3, 12.00%) |
| Ether lipid metabolism(3, 18.75%) |
| Glycosaminoglycan degradation(3, 37.50%) |
| Methane metabolism(3, 18.75%) |
| Cholesterol metabolism(3, 23.08%) |
| Toll and Imd signaling pathway(2, 6.90%) |
| Platelet activation(2, 4.35%) |
| Circadian entrainment(2, 5.41%) |
| Serotonergic synapse(2, 5.56%) |
| Inflammatory mediator regulation of TRP channels(2, 6.06%) |
| Aldosterone synthesis and secretion(2, 5.56%) |
| Salivary secretion(2, 5.56%) |
| NF-kappa B signaling pathway(2, 7.69%) |
| Toll-like receptor signaling pathway(2, 6.90%) |
| Leishmaniasis(2, 9.52%) |
| Chagas disease(2, 5.71%) |
| Mitophagy - animal(2, 5.41%) |
| Central carbon metabolism in cancer(2, 6.67%) |
| Longevity regulating pathway(2, 4.17%) |
| Mannose type O-glycan biosynthesis(2, 25.00%) |
| Olfactory transduction(2, 11.76%) |
| Morphine addiction(2, 6.25%) |
| Ribosome biogenesis in eukaryotes(2, 3.12%) |
| Systemic lupus erythematosus(2, 12.50%) |
| Cardiac muscle contraction(2, 7.69%) |
| Arrhythmogenic right ventricular cardiomyopathy(2, 10.00%) |
| Dilated cardiomyopathy(2, 6.90%) |
| Cytosolic DNA-sensing pathway(2, 9.09%) |
| VEGF signaling pathway(2, 9.09%) |
| Leukocyte transendothelial migration(2, 7.14%) |
| N-Glycan biosynthesis(2, 5.71%) |
| Nucleotide excision repair(2, 5.71%) |
| Base excision repair(2, 8.33%) |
| B cell receptor signaling pathway(2, 8.70%) |
| Chronic myeloid leukemia(2, 6.25%) |
| Butanoate metabolism(2, 14.29%) |
| DNA replication(2, 6.25%) |
| Apoptosis - multiple species(2, 12.50%) |
| Cell adhesion molecules(2, 10.00%) |
| Mitophagy - yeast(2, 11.76%) |
| Vasopressin-regulated water reabsorption(2, 8.70%) |
| Other glycan degradation(2, 20.00%) |
| Glycosphingolipid biosynthesis - globo and isoglobo series(2, 33.33%) |
| Endometrial cancer(2, 7.69%) |
| Pyrimidine metabolism(2, 6.90%) |
| Sphingolipid signaling pathway(2, 4.55%) |
| Prolactin signaling pathway(2, 7.41%) |
| Cocaine addiction(2, 9.09%) |
| Arginine and proline metabolism(2, 8.33%) |
| Mucin type O-glycan biosynthesis(2, 50.00%) |
| Insulin secretion(2, 6.06%) |
| Bile secretion(2, 8.70%) |
| Quorum sensing(2, 40.00%) |
| Selenocompound metabolism(2, 22.22%) |
| Renin-angiotensin system(2, 28.57%) |
| Phenylalanine metabolism(2, 20.00%) |
| Isoquinoline alkaloid biosynthesis(2, 25.00%) |
| Ascorbate and aldarate metabolism(2, 22.22%) |
| Propanoate metabolism(2, 8.70%) |
| Arachidonic acid metabolism(2, 12.50%) |
| Glycosaminoglycan biosynthesis - keratan sulfate(2, 25.00%) |
| Endocrine and other factor-regulated calcium reabsorption(2, 10.00%) |
| alpha-Linolenic acid metabolism(2, 50.00%) |
| Mineral absorption(2, 12.50%) |
| Chloroalkane and chloroalkene degradation(2, 66.67%) |
| Histidine metabolism(2, 16.67%) |
| Phosphonate and phosphinate metabolism(2, 66.67%) |
| Apelin signaling pathway(1, 1.89%) |
| C-type lectin receptor signaling pathway(1, 3.12%) |
| Long-term potentiation(1, 4.00%) |
| Glutamatergic synapse(1, 2.38%) |
| Long-term depression(1, 4.55%) |
| Phototransduction - fly(1, 5.88%) |
| Renin secretion(1, 3.70%) |
| Cortisol synthesis and secretion(1, 4.17%) |
| GnRH secretion(1, 5.26%) |
| Gastric acid secretion(1, 3.85%) |
| Autophagy - other(1, 4.76%) |
| Autophagy - yeast(1, 1.89%) |
| Terpenoid backbone biosynthesis(1, 5.26%) |
| Pertussis(1, 4.00%) |
| Renal cell carcinoma(1, 3.23%) |
| Pyruvate metabolism(1, 4.17%) |
| Type II diabetes mellitus(1, 5.56%) |
| Hedgehog signaling pathway(1, 4.35%) |
| Hedgehog signaling pathway - fly(1, 4.76%) |
| TGF-beta signaling pathway(1, 3.03%) |
| Basal transcription factors(1, 3.23%) |
| Longevity regulating pathway - multiple species(1, 2.63%) |
| Fatty acid elongation(1, 7.69%) |
| Steroid hormone biosynthesis(1, 11.11%) |
| Staphylococcus aureus infection(1, 14.29%) |
| Nicotine addiction(1, 10.00%) |
| Circadian rhythm(1, 7.69%) |
| Pantothenate and CoA biosynthesis(1, 9.09%) |
| RNA degradation(1, 1.79%) |
| Malaria(1, 9.09%) |
| ABC transporters(1, 5.88%) |
| Mismatch repair(1, 5.00%) |
| Thyroid cancer(1, 7.14%) |
| Vitamin digestion and absorption(1, 12.50%) |
| Basal cell carcinoma(1, 3.45%) |
| Antifolate resistance(1, 7.69%) |
| O-Antigen nucleotide sugar biosynthesis(1, 16.67%) |
| Non-homologous end-joining(1, 10.00%) |
| Melanoma(1, 5.00%) |
| Bladder cancer(1, 7.69%) |
| Nitrogen metabolism(1, 16.67%) |
| Bacterial secretion system(1, 33.33%) |
| MAPK signaling pathway - plant(1, 10.00%) |
| Cutin, suberine and wax biosynthesis(1, 100.00%) |
| Hematopoietic cell lineage(1, 12.50%) |
| Fatty acid biosynthesis(1, 12.50%) |
| Ferroptosis(1, 6.67%) |
| Indole alkaloid biosynthesis(1, 100.00%) |
| Betalain biosynthesis(1, 100.00%) |
| Nicotinate and nicotinamide metabolism(1, 6.67%) |
| Thiamine metabolism(1, 20.00%) |
| Pancreatic cancer(1, 3.23%) |
| Circadian rhythm - fly(1, 14.29%) |
| Pentose phosphate pathway(1, 6.25%) |
| Glycosphingolipid biosynthesis - ganglio series(1, 25.00%) |
| African trypanosomiasis(1, 20.00%) |
| SNARE interactions in vesicular transport(1, 4.76%) |
| Linoleic acid metabolism(1, 20.00%) |
| Degradation of aromatic compounds(1, 33.33%) |
| Naphthalene degradation(1, 50.00%) |
| Steroid biosynthesis(1, 12.50%) |
| Two-component system(1, 7.69%) |
| Limonene and pinene degradation(1, 100.00%) |
| Aldosterone-regulated sodium reabsorption(1, 8.33%) |
| Primary bile acid biosynthesis(1, 25.00%) |
| Streptomycin biosynthesis(1, 25.00%) |
| Neomycin, kanamycin and gentamicin biosynthesis(1, 100.00%) |
| Cyanoamino acid metabolism(1, 25.00%) |
| Plant hormone signal transduction(1, 33.33%) |
| Styrene degradation(1, 50.00%) |
| Biofilm formation - Escherichia coli(1, 50.00%) |

Pathway Sorted by Name

| Pathway  Pathway |
| --- |
| ABC transporters(1, 5.88%) |
| AGE-RAGE signaling pathway in diabetic complications(4, 11.76%) |
| AMPK signaling pathway(5, 9.09%) |
| Acute myeloid leukemia(3, 13.64%) |
| Adherens junction(3, 8.33%) |
| Adipocytokine signaling pathway(5, 20.00%) |
| Adrenergic signaling in cardiomyocytes(3, 6.98%) |
| African trypanosomiasis(1, 20.00%) |
| Alanine, aspartate and glutamate metabolism(3, 12.50%) |
| Alcoholism(5, 10.64%) |
| Aldosterone synthesis and secretion(2, 5.56%) |
| Aldosterone-regulated sodium reabsorption(1, 8.33%) |
| Amino sugar and nucleotide sugar metabolism(5, 16.67%) |
| Aminoacyl-tRNA biosynthesis(3, 11.54%) |
| Amoebiasis(3, 10.00%) |
| Amphetamine addiction(3, 10.71%) |
| Antifolate resistance(1, 7.69%) |
| Antigen processing and presentation(6, 37.50%) |
| Apelin signaling pathway(1, 1.89%) |
| Apoptosis(10, 17.86%) |
| Apoptosis - fly(5, 12.20%) |
| Apoptosis - multiple species(2, 12.50%) |
| Arachidonic acid metabolism(2, 12.50%) |
| Arginine and proline metabolism(2, 8.33%) |
| Arrhythmogenic right ventricular cardiomyopathy(2, 10.00%) |
| Ascorbate and aldarate metabolism(2, 22.22%) |
| Autophagy - animal(5, 6.25%) |
| Autophagy - other(1, 4.76%) |
| Autophagy - yeast(1, 1.89%) |
| Axon guidance(3, 4.55%) |
| Axon regeneration(3, 6.25%) |
| B cell receptor signaling pathway(2, 8.70%) |
| Bacterial secretion system(1, 33.33%) |
| Basal cell carcinoma(1, 3.45%) |
| Basal transcription factors(1, 3.23%) |
| Base excision repair(2, 8.33%) |
| Betalain biosynthesis(1, 100.00%) |
| Bile secretion(2, 8.70%) |
| Biofilm formation - Escherichia coli(1, 50.00%) |
| Biosynthesis of amino acids(4, 9.52%) |
| Biosynthesis of cofactors(15, 18.29%) |
| Biosynthesis of unsaturated fatty acids(4, 36.36%) |
| Bladder cancer(1, 7.69%) |
| Breast cancer(4, 7.14%) |
| Butanoate metabolism(2, 14.29%) |
| C-type lectin receptor signaling pathway(1, 3.12%) |
| Calcium signaling pathway(8, 11.27%) |
| Carbohydrate digestion and absorption(3, 21.43%) |
| Carbon metabolism(7, 9.59%) |
| Cardiac muscle contraction(2, 7.69%) |
| Cell adhesion molecules(2, 10.00%) |
| Cell cycle(14, 19.18%) |
| Cell cycle - yeast(9, 14.75%) |
| Cellular senescence(5, 7.46%) |
| Central carbon metabolism in cancer(2, 6.67%) |
| Chagas disease(2, 5.71%) |
| Chemical carcinogenesis(5, 33.33%) |
| Chemokine signaling pathway(5, 8.93%) |
| Chloroalkane and chloroalkene degradation(2, 66.67%) |
| Cholesterol metabolism(3, 23.08%) |
| Choline metabolism in cancer(7, 18.92%) |
| Cholinergic synapse(3, 7.14%) |
| Chronic myeloid leukemia(2, 6.25%) |
| Circadian entrainment(2, 5.41%) |
| Circadian rhythm(1, 7.69%) |
| Circadian rhythm - fly(1, 14.29%) |
| Cocaine addiction(2, 9.09%) |
| Colorectal cancer(3, 7.89%) |
| Complement and coagulation cascades(3, 21.43%) |
| Cortisol synthesis and secretion(1, 4.17%) |
| Cushing syndrome(6, 8.70%) |
| Cutin, suberine and wax biosynthesis(1, 100.00%) |
| Cyanoamino acid metabolism(1, 25.00%) |
| Cysteine and methionine metabolism(5, 16.13%) |
| Cytosolic DNA-sensing pathway(2, 9.09%) |
| DNA replication(2, 6.25%) |
| Degradation of aromatic compounds(1, 33.33%) |
| Dilated cardiomyopathy(2, 6.90%) |
| Dopaminergic synapse(3, 6.67%) |
| Dorso-ventral axis formation(5, 22.73%) |
| Drug metabolism - cytochrome P450(5, 55.56%) |
| Drug metabolism - other enzymes(4, 18.18%) |
| ECM-receptor interaction(6, 27.27%) |
| EGFR tyrosine kinase inhibitor resistance(5, 17.24%) |
| Endocrine and other factor-regulated calcium reabsorption(2, 10.00%) |
| Endocrine resistance(3, 6.82%) |
| Endometrial cancer(2, 7.69%) |
| Epithelial cell signaling in Helicobacter pylori infection(3, 9.38%) |
| Epstein-Barr virus infection(7, 9.86%) |
| ErbB signaling pathway(4, 12.12%) |
| Estrogen signaling pathway(9, 18.75%) |
| Ether lipid metabolism(3, 18.75%) |
| Fat digestion and absorption(3, 23.08%) |
| Fatty acid biosynthesis(1, 12.50%) |
| Fatty acid degradation(4, 15.38%) |
| Fatty acid elongation(1, 7.69%) |
| Fatty acid metabolism(5, 15.15%) |
| Fc epsilon RI signaling pathway(3, 14.29%) |
| Fc gamma R-mediated phagocytosis(3, 8.33%) |
| Ferroptosis(1, 6.67%) |
| Fluid shear stress and atherosclerosis(4, 9.09%) |
| Focal adhesion(9, 11.54%) |
| Folate biosynthesis(5, 29.41%) |
| FoxO signaling pathway(7, 12.50%) |
| Fructose and mannose metabolism(6, 31.58%) |
| GABAergic synapse(3, 9.68%) |
| Galactose metabolism(4, 22.22%) |
| Gap junction(3, 8.82%) |
| Gastric acid secretion(1, 3.85%) |
| Gastric cancer(5, 9.09%) |
| Glioma(4, 14.81%) |
| Glucagon signaling pathway(3, 6.98%) |
| Glutamatergic synapse(1, 2.38%) |
| Glutathione metabolism(4, 16.67%) |
| Glycerolipid metabolism(4, 14.81%) |
| Glycerophospholipid metabolism(4, 10.00%) |
| Glycine, serine and threonine metabolism(5, 17.86%) |
| Glycolysis / Gluconeogenesis(7, 21.88%) |
| Glycosaminoglycan biosynthesis - keratan sulfate(2, 25.00%) |
| Glycosaminoglycan degradation(3, 37.50%) |
| Glycosphingolipid biosynthesis - ganglio series(1, 25.00%) |
| Glycosphingolipid biosynthesis - globo and isoglobo series(2, 33.33%) |
| Glycosphingolipid biosynthesis - lacto and neolacto series(5, 45.45%) |
| Glyoxylate and dicarboxylate metabolism(3, 12.00%) |
| GnRH secretion(1, 5.26%) |
| GnRH signaling pathway(4, 10.00%) |
| Growth hormone synthesis, secretion and action(5, 10.00%) |
| HIF-1 signaling pathway(5, 12.20%) |
| Hedgehog signaling pathway(1, 4.35%) |
| Hedgehog signaling pathway - fly(1, 4.76%) |
| Hematopoietic cell lineage(1, 12.50%) |
| Hepatitis B(9, 15.52%) |
| Hepatitis C(5, 12.50%) |
| Hepatocellular carcinoma(7, 10.45%) |
| Herpes simplex virus 1 infection(4, 7.27%) |
| Hippo signaling pathway(3, 5.26%) |
| Hippo signaling pathway - fly(6, 13.64%) |
| Histidine metabolism(2, 16.67%) |
| Human T-cell leukemia virus 1 infection(10, 11.11%) |
| Human cytomegalovirus infection(5, 6.58%) |
| Human immunodeficiency virus 1 infection(4, 5.06%) |
| Hypertrophic cardiomyopathy(3, 12.50%) |
| IL-17 signaling pathway(3, 11.11%) |
| Indole alkaloid biosynthesis(1, 100.00%) |
| Inflammatory mediator regulation of TRP channels(2, 6.06%) |
| Influenza A(6, 13.64%) |
| Inositol phosphate metabolism(3, 8.33%) |
| Insect hormone biosynthesis(3, 30.00%) |
| Insulin resistance(4, 8.89%) |
| Insulin secretion(2, 6.06%) |
| Insulin signaling pathway(3, 5.17%) |
| Isoquinoline alkaloid biosynthesis(2, 25.00%) |
| JAK-STAT signaling pathway(3, 14.29%) |
| Kaposi sarcoma-associated herpesvirus infection(3, 5.77%) |
| Legionellosis(5, 21.74%) |
| Leishmaniasis(2, 9.52%) |
| Leukocyte transendothelial migration(2, 7.14%) |
| Limonene and pinene degradation(1, 100.00%) |
| Linoleic acid metabolism(1, 20.00%) |
| Long-term depression(1, 4.55%) |
| Long-term potentiation(1, 4.00%) |
| Longevity regulating pathway(2, 4.17%) |
| Longevity regulating pathway - multiple species(1, 2.63%) |
| Longevity regulating pathway - worm(3, 7.69%) |
| Lysine degradation(5, 13.89%) |
| Lysosome(11, 17.46%) |
| MAPK signaling pathway(5, 5.56%) |
| MAPK signaling pathway - fly(4, 6.06%) |
| MAPK signaling pathway - plant(1, 10.00%) |
| MAPK signaling pathway - yeast(3, 10.34%) |
| Malaria(1, 9.09%) |
| Mannose type O-glycan biosynthesis(2, 25.00%) |
| Measles(6, 13.64%) |
| Meiosis - yeast(7, 14.89%) |
| Melanogenesis(3, 7.50%) |
| Melanoma(1, 5.00%) |
| Metabolism of xenobiotics by cytochrome P450(5, 55.56%) |
| Methane metabolism(3, 18.75%) |
| MicroRNAs in cancer(8, 13.33%) |
| Mineral absorption(2, 12.50%) |
| Mismatch repair(1, 5.00%) |
| Mitophagy - animal(2, 5.41%) |
| Mitophagy - yeast(2, 11.76%) |
| Morphine addiction(2, 6.25%) |
| Mucin type O-glycan biosynthesis(2, 50.00%) |
| N-Glycan biosynthesis(2, 5.71%) |
| NF-kappa B signaling pathway(2, 7.69%) |
| NOD-like receptor signaling pathway(5, 10.00%) |
| Naphthalene degradation(1, 50.00%) |
| Natural killer cell mediated cytotoxicity(3, 13.64%) |
| Necroptosis(4, 11.11%) |
| Neomycin, kanamycin and gentamicin biosynthesis(1, 100.00%) |
| Neuroactive ligand-receptor interaction(4, 6.67%) |
| Neurotrophin signaling pathway(3, 5.88%) |
| Nicotinate and nicotinamide metabolism(1, 6.67%) |
| Nicotine addiction(1, 10.00%) |
| Nitrogen metabolism(1, 16.67%) |
| Non-alcoholic fatty liver disease(4, 4.76%) |
| Non-homologous end-joining(1, 10.00%) |
| Non-small cell lung cancer(4, 16.00%) |
| Notch signaling pathway(3, 12.00%) |
| Nucleotide excision repair(2, 5.71%) |
| O-Antigen nucleotide sugar biosynthesis(1, 16.67%) |
| Olfactory transduction(2, 11.76%) |
| One carbon pool by folate(5, 45.45%) |
| Oocyte meiosis(10, 16.67%) |
| Osteoclast differentiation(3, 10.71%) |
| Other glycan degradation(2, 20.00%) |
| Other types of O-glycan biosynthesis(5, 33.33%) |
| Oxytocin signaling pathway(3, 5.26%) |
| PD-L1 expression and PD-1 checkpoint pathway in cancer(4, 12.90%) |
| PI3K-Akt signaling pathway(15, 14.56%) |
| PPAR signaling pathway(4, 14.29%) |
| Pancreatic cancer(1, 3.23%) |
| Pancreatic secretion(5, 12.20%) |
| Pantothenate and CoA biosynthesis(1, 9.09%) |
| Parathyroid hormone synthesis, secretion and action(5, 11.11%) |
| Pathogenic Escherichia coli infection(6, 9.38%) |
| Pentose and glucuronate interconversions(3, 21.43%) |
| Pentose phosphate pathway(1, 6.25%) |
| Peroxisome(10, 21.74%) |
| Pertussis(1, 4.00%) |
| Phagosome(5, 9.80%) |
| Phenylalanine metabolism(2, 20.00%) |
| Phosphatidylinositol signaling system(3, 7.14%) |
| Phospholipase D signaling pathway(4, 7.55%) |
| Phosphonate and phosphinate metabolism(2, 66.67%) |
| Phototransduction - fly(1, 5.88%) |
| Plant hormone signal transduction(1, 33.33%) |
| Plant-pathogen interaction(3, 27.27%) |
| Platelet activation(2, 4.35%) |
| Platinum drug resistance(4, 14.29%) |
| Porphyrin and chlorophyll metabolism(6, 35.29%) |
| Primary bile acid biosynthesis(1, 25.00%) |
| Progesterone-mediated oocyte maturation(8, 17.02%) |
| Prolactin signaling pathway(2, 7.41%) |
| Propanoate metabolism(2, 8.70%) |
| Prostate cancer(8, 20.51%) |
| Protein digestion and absorption(8, 24.24%) |
| Protein export(7, 33.33%) |
| Proteoglycans in cancer(9, 9.68%) |
| Purine metabolism(4, 5.97%) |
| Pyrimidine metabolism(2, 6.90%) |
| Pyruvate metabolism(1, 4.17%) |
| Quorum sensing(2, 40.00%) |
| RIG-I-like receptor signaling pathway(3, 17.65%) |
| RNA degradation(1, 1.79%) |
| Rap1 signaling pathway(5, 6.76%) |
| Ras signaling pathway(6, 7.79%) |
| Regulation of actin cytoskeleton(5, 6.85%) |
| Relaxin signaling pathway(7, 12.73%) |
| Renal cell carcinoma(1, 3.23%) |
| Renin secretion(1, 3.70%) |
| Renin-angiotensin system(2, 28.57%) |
| Retinol metabolism(4, 36.36%) |
| Retrograde endocannabinoid signaling(3, 4.41%) |
| Rheumatoid arthritis(3, 14.29%) |
| Ribosome biogenesis in eukaryotes(2, 3.12%) |
| SNARE interactions in vesicular transport(1, 4.76%) |
| Salivary secretion(2, 5.56%) |
| Salmonella infection(5, 4.95%) |
| Selenocompound metabolism(2, 22.22%) |
| Serotonergic synapse(2, 5.56%) |
| Shigellosis(4, 3.88%) |
| Signaling pathways regulating pluripotency of stem cells(6, 11.54%) |
| Small cell lung cancer(6, 15.79%) |
| Sphingolipid signaling pathway(2, 4.55%) |
| Spinocerebellar ataxia(5, 6.02%) |
| Staphylococcus aureus infection(1, 14.29%) |
| Starch and sucrose metabolism(5, 23.81%) |
| Steroid biosynthesis(1, 12.50%) |
| Steroid hormone biosynthesis(1, 11.11%) |
| Streptomycin biosynthesis(1, 25.00%) |
| Styrene degradation(1, 50.00%) |
| Systemic lupus erythematosus(2, 12.50%) |
| T cell receptor signaling pathway(3, 8.33%) |
| TGF-beta signaling pathway(1, 3.03%) |
| TNF signaling pathway(4, 10.81%) |
| Terpenoid backbone biosynthesis(1, 5.26%) |
| Th1 and Th2 cell differentiation(4, 17.39%) |
| Th17 cell differentiation(4, 16.00%) |
| Thiamine metabolism(1, 20.00%) |
| Thyroid cancer(1, 7.14%) |
| Thyroid hormone signaling pathway(4, 7.02%) |
| Thyroid hormone synthesis(6, 18.18%) |
| Tight junction(5, 7.58%) |
| Toll and Imd signaling pathway(2, 6.90%) |
| Toll-like receptor signaling pathway(2, 6.90%) |
| Toxoplasmosis(4, 11.11%) |
| Transcriptional misregulation in cancer(9, 12.50%) |
| Tryptophan metabolism(4, 16.00%) |
| Tuberculosis(4, 8.00%) |
| Two-component system(1, 7.69%) |
| Type II diabetes mellitus(1, 5.56%) |
| Tyrosine metabolism(5, 27.78%) |
| Ubiquitin mediated proteolysis(6, 7.32%) |
| VEGF signaling pathway(2, 9.09%) |
| Valine, leucine and isoleucine degradation(3, 8.57%) |
| Various types of N-glycan biosynthesis(4, 15.38%) |
| Vascular smooth muscle contraction(4, 9.30%) |
| Vasopressin-regulated water reabsorption(2, 8.70%) |
| Vibrio cholerae infection(6, 20.00%) |
| Viral carcinogenesis(11, 13.92%) |
| Vitamin digestion and absorption(1, 12.50%) |
| Wnt signaling pathway(5, 6.94%) |
| Yersinia infection(3, 4.92%) |
| alpha-Linolenic acid metabolism(2, 50.00%) |
| beta-Alanine metabolism(4, 23.53%) |
| cAMP signaling pathway(4, 5.88%) |
| cGMP-PKG signaling pathway(4, 6.78%) |
| mRNA surveillance pathway(4, 7.02%) |
| mTOR signaling pathway(5, 5.75%) |
| p53 signaling pathway(6, 20.00%) |
