## Supplementary material for "The transcriptomic signature of obligate parthenogenesis": Supplementary File 6-select_UD(red)_OD(green).html

  

Pathway Sorted by Number of Associated Genes

| Pathway  Pathway |
| --- |
| Cell cycle(10, 13.70%) |
| Viral carcinogenesis(9, 11.39%) |
| Human T-cell leukemia virus 1 infection(8, 8.89%) |
| Biosynthesis of cofactors(8, 9.76%) |
| Cell cycle - yeast(8, 13.11%) |
| Peroxisome(8, 17.39%) |
| Hepatitis B(7, 12.07%) |
| Carbon metabolism(7, 9.59%) |
| PI3K-Akt signaling pathway(7, 6.80%) |
| Apoptosis(6, 10.71%) |
| Influenza A(6, 13.64%) |
| Ubiquitin mediated proteolysis(6, 7.32%) |
| Meiosis - yeast(6, 12.77%) |
| Lysosome(6, 9.52%) |
| Calcium signaling pathway(5, 7.04%) |
| Oocyte meiosis(5, 8.33%) |
| Estrogen signaling pathway(5, 10.42%) |
| Thyroid hormone synthesis(5, 15.15%) |
| Proteoglycans in cancer(5, 5.38%) |
| Choline metabolism in cancer(5, 13.51%) |
| Prostate cancer(5, 12.82%) |
| Alcoholism(5, 10.64%) |
| One carbon pool by folate(5, 45.45%) |
| Hepatocellular carcinoma(5, 7.46%) |
| Amino sugar and nucleotide sugar metabolism(5, 16.67%) |
| Small cell lung cancer(5, 13.16%) |
| Fructose and mannose metabolism(5, 26.32%) |
| Protein export(5, 23.81%) |
| Signaling pathways regulating pluripotency of stem cells(5, 9.62%) |
| p53 signaling pathway(5, 16.67%) |
| Cushing syndrome(4, 5.80%) |
| cGMP-PKG signaling pathway(4, 6.78%) |
| Growth hormone synthesis, secretion and action(4, 8.00%) |
| Human cytomegalovirus infection(4, 5.26%) |
| Pathogenic Escherichia coli infection(4, 6.25%) |
| Measles(4, 9.09%) |
| Herpes simplex virus 1 infection(4, 7.27%) |
| Epstein-Barr virus infection(4, 5.63%) |
| Tight junction(4, 6.06%) |
| Progesterone-mediated oocyte maturation(4, 8.51%) |
| Biosynthesis of unsaturated fatty acids(4, 36.36%) |
| Fatty acid metabolism(4, 12.12%) |
| Chemokine signaling pathway(4, 7.14%) |
| cAMP signaling pathway(4, 5.88%) |
| FoxO signaling pathway(4, 7.14%) |
| MicroRNAs in cancer(4, 6.67%) |
| Folate biosynthesis(4, 23.53%) |
| Hepatitis C(4, 10.00%) |
| Relaxin signaling pathway(4, 7.27%) |
| Focal adhesion(4, 5.13%) |
| mTOR signaling pathway(4, 4.60%) |
| Platinum drug resistance(4, 14.29%) |
| Protein digestion and absorption(4, 12.12%) |
| Glutathione metabolism(4, 16.67%) |
| Metabolism of xenobiotics by cytochrome P450(4, 44.44%) |
| Drug metabolism - cytochrome P450(4, 44.44%) |
| Chemical carcinogenesis(4, 26.67%) |
| Cysteine and methionine metabolism(4, 12.90%) |
| Biosynthesis of amino acids(4, 9.52%) |
| Glycine, serine and threonine metabolism(4, 14.29%) |
| Glycolysis / Gluconeogenesis(4, 12.50%) |
| Autophagy - animal(3, 3.75%) |
| Cellular senescence(3, 4.48%) |
| Vascular smooth muscle contraction(3, 6.98%) |
| NOD-like receptor signaling pathway(3, 6.00%) |
| Cholinergic synapse(3, 7.14%) |
| Dopaminergic synapse(3, 6.67%) |
| GnRH signaling pathway(3, 7.50%) |
| Parathyroid hormone synthesis, secretion and action(3, 6.67%) |
| Spinocerebellar ataxia(3, 3.61%) |
| Human immunodeficiency virus 1 infection(3, 3.80%) |
| Salmonella infection(3, 2.97%) |
| Toxoplasmosis(3, 8.33%) |
| Tuberculosis(3, 6.00%) |
| Axon regeneration(3, 6.25%) |
| Lysine degradation(3, 8.33%) |
| Regulation of actin cytoskeleton(3, 4.11%) |
| Purine metabolism(3, 4.48%) |
| PPAR signaling pathway(3, 10.71%) |
| AMPK signaling pathway(3, 5.45%) |
| Adipocytokine signaling pathway(3, 12.00%) |
| Necroptosis(3, 8.33%) |
| Fluid shear stress and atherosclerosis(3, 6.82%) |
| Transcriptional misregulation in cancer(3, 4.17%) |
| Wnt signaling pathway(3, 4.17%) |
| Melanogenesis(3, 7.50%) |
| TNF signaling pathway(3, 8.11%) |
| Adrenergic signaling in cardiomyocytes(3, 6.98%) |
| EGFR tyrosine kinase inhibitor resistance(3, 10.34%) |
| Insulin resistance(3, 6.67%) |
| Legionellosis(3, 13.04%) |
| Fatty acid degradation(3, 11.54%) |
| Glycerolipid metabolism(3, 11.11%) |
| Gastric cancer(3, 5.45%) |
| Insect hormone biosynthesis(3, 30.00%) |
| Tryptophan metabolism(3, 12.00%) |
| Amphetamine addiction(3, 10.71%) |
| Glyoxylate and dicarboxylate metabolism(3, 12.00%) |
| Tyrosine metabolism(3, 16.67%) |
| Methane metabolism(3, 18.75%) |
| beta-Alanine metabolism(3, 17.65%) |
| Phosphatidylinositol signaling system(2, 4.76%) |
| Gap junction(2, 5.88%) |
| Retrograde endocannabinoid signaling(2, 2.94%) |
| Serotonergic synapse(2, 5.56%) |
| Oxytocin signaling pathway(2, 3.51%) |
| Glucagon signaling pathway(2, 4.65%) |
| Aldosterone synthesis and secretion(2, 5.56%) |
| Pancreatic secretion(2, 4.88%) |
| Shigellosis(2, 1.94%) |
| Kaposi sarcoma-associated herpesvirus infection(2, 3.85%) |
| MAPK signaling pathway(2, 2.22%) |
| Ras signaling pathway(2, 2.60%) |
| T cell receptor signaling pathway(2, 5.56%) |
| PD-L1 expression and PD-1 checkpoint pathway in cancer(2, 6.45%) |
| Neurotrophin signaling pathway(2, 3.92%) |
| Yersinia infection(2, 3.28%) |
| Leishmaniasis(2, 9.52%) |
| HIF-1 signaling pathway(2, 4.88%) |
| Longevity regulating pathway - worm(2, 5.13%) |
| Th17 cell differentiation(2, 8.00%) |
| Thyroid hormone signaling pathway(2, 3.51%) |
| Fc gamma R-mediated phagocytosis(2, 5.56%) |
| Longevity regulating pathway(2, 4.17%) |
| Non-alcoholic fatty liver disease(2, 2.38%) |
| Antigen processing and presentation(2, 12.50%) |
| Plant-pathogen interaction(2, 18.18%) |
| IL-17 signaling pathway(2, 7.41%) |
| Adherens junction(2, 5.56%) |
| JAK-STAT signaling pathway(2, 9.52%) |
| Ribosome biogenesis in eukaryotes(2, 3.12%) |
| Aminoacyl-tRNA biosynthesis(2, 7.69%) |
| Osteoclast differentiation(2, 7.14%) |
| Carbohydrate digestion and absorption(2, 14.29%) |
| MAPK signaling pathway - yeast(2, 6.90%) |
| Systemic lupus erythematosus(2, 12.50%) |
| Cardiac muscle contraction(2, 7.69%) |
| Hypertrophic cardiomyopathy(2, 8.33%) |
| Arrhythmogenic right ventricular cardiomyopathy(2, 10.00%) |
| Dilated cardiomyopathy(2, 6.90%) |
| Cytosolic DNA-sensing pathway(2, 9.09%) |
| ErbB signaling pathway(2, 6.06%) |
| Rap1 signaling pathway(2, 2.70%) |
| Phospholipase D signaling pathway(2, 3.77%) |
| Natural killer cell mediated cytotoxicity(2, 9.09%) |
| Th1 and Th2 cell differentiation(2, 8.70%) |
| Fc epsilon RI signaling pathway(2, 9.52%) |
| AGE-RAGE signaling pathway in diabetic complications(2, 5.88%) |
| Glioma(2, 7.41%) |
| Non-small cell lung cancer(2, 8.00%) |
| Phagosome(2, 3.92%) |
| B cell receptor signaling pathway(2, 8.70%) |
| RIG-I-like receptor signaling pathway(2, 11.76%) |
| Chronic myeloid leukemia(2, 6.25%) |
| ECM-receptor interaction(2, 9.09%) |
| Various types of N-glycan biosynthesis(2, 7.69%) |
| Apoptosis - fly(2, 4.88%) |
| Cell adhesion molecules(2, 10.00%) |
| Vasopressin-regulated water reabsorption(2, 8.70%) |
| Breast cancer(2, 3.57%) |
| Hippo signaling pathway(2, 3.51%) |
| Hippo signaling pathway - fly(2, 4.55%) |
| Drug metabolism - other enzymes(2, 9.09%) |
| Glycerophospholipid metabolism(2, 5.00%) |
| Insulin signaling pathway(2, 3.45%) |
| Other glycan degradation(2, 20.00%) |
| Amoebiasis(2, 6.67%) |
| Glycosphingolipid biosynthesis - lacto and neolacto series(2, 18.18%) |
| Glycosphingolipid biosynthesis - globo and isoglobo series(2, 33.33%) |
| Sphingolipid signaling pathway(2, 4.55%) |
| Dorso-ventral axis formation(2, 9.09%) |
| Prolactin signaling pathway(2, 7.41%) |
| Colorectal cancer(2, 5.26%) |
| Cocaine addiction(2, 9.09%) |
| Arginine and proline metabolism(2, 8.33%) |
| Other types of O-glycan biosynthesis(2, 13.33%) |
| Insulin secretion(2, 6.06%) |
| Ether lipid metabolism(2, 12.50%) |
| Selenocompound metabolism(2, 22.22%) |
| Phenylalanine metabolism(2, 20.00%) |
| Retinol metabolism(2, 18.18%) |
| Porphyrin and chlorophyll metabolism(2, 11.76%) |
| Endocrine and other factor-regulated calcium reabsorption(2, 10.00%) |
| Cholesterol metabolism(2, 15.38%) |
| Mineral absorption(2, 12.50%) |
| Chloroalkane and chloroalkene degradation(2, 66.67%) |
| Histidine metabolism(2, 16.67%) |
| Phosphonate and phosphinate metabolism(2, 66.67%) |
| Toll and Imd signaling pathway(1, 3.45%) |
| Apelin signaling pathway(1, 1.89%) |
| Platelet activation(1, 2.17%) |
| C-type lectin receptor signaling pathway(1, 3.12%) |
| Circadian entrainment(1, 2.70%) |
| Long-term potentiation(1, 4.00%) |
| Glutamatergic synapse(1, 2.38%) |
| Long-term depression(1, 4.55%) |
| Phototransduction - fly(1, 5.88%) |
| Inflammatory mediator regulation of TRP channels(1, 3.03%) |
| Cortisol synthesis and secretion(1, 4.17%) |
| GnRH secretion(1, 5.26%) |
| Salivary secretion(1, 2.78%) |
| Gastric acid secretion(1, 3.85%) |
| Notch signaling pathway(1, 4.00%) |
| Inositol phosphate metabolism(1, 2.78%) |
| NF-kappa B signaling pathway(1, 3.85%) |
| Toll-like receptor signaling pathway(1, 3.45%) |
| Chagas disease(1, 2.86%) |
| Mitophagy - animal(1, 2.70%) |
| Renal cell carcinoma(1, 3.23%) |
| Central carbon metabolism in cancer(1, 3.33%) |
| Pyruvate metabolism(1, 4.17%) |
| Type II diabetes mellitus(1, 5.56%) |
| Hedgehog signaling pathway(1, 4.35%) |
| Hedgehog signaling pathway - fly(1, 4.76%) |
| Olfactory transduction(1, 5.88%) |
| Morphine addiction(1, 3.12%) |
| mRNA surveillance pathway(1, 1.75%) |
| TGF-beta signaling pathway(1, 3.03%) |
| Basal transcription factors(1, 3.23%) |
| MAPK signaling pathway - fly(1, 1.52%) |
| Galactose metabolism(1, 5.56%) |
| Fatty acid elongation(1, 7.69%) |
| Axon guidance(1, 1.52%) |
| VEGF signaling pathway(1, 4.55%) |
| Leukocyte transendothelial migration(1, 3.57%) |
| Vibrio cholerae infection(1, 3.33%) |
| Epithelial cell signaling in Helicobacter pylori infection(1, 3.12%) |
| Complement and coagulation cascades(1, 7.14%) |
| Neuroactive ligand-receptor interaction(1, 1.67%) |
| GABAergic synapse(1, 3.23%) |
| N-Glycan biosynthesis(1, 2.86%) |
| Nucleotide excision repair(1, 2.86%) |
| Pantothenate and CoA biosynthesis(1, 9.09%) |
| Base excision repair(1, 4.17%) |
| Malaria(1, 9.09%) |
| ABC transporters(1, 5.88%) |
| Valine, leucine and isoleucine degradation(1, 2.86%) |
| Butanoate metabolism(1, 7.14%) |
| DNA replication(1, 3.12%) |
| Mismatch repair(1, 5.00%) |
| Apoptosis - multiple species(1, 6.25%) |
| Mitophagy - yeast(1, 5.88%) |
| Fat digestion and absorption(1, 7.69%) |
| Vitamin digestion and absorption(1, 12.50%) |
| Basal cell carcinoma(1, 3.45%) |
| Antifolate resistance(1, 7.69%) |
| Pentose and glucuronate interconversions(1, 7.14%) |
| O-Antigen nucleotide sugar biosynthesis(1, 16.67%) |
| Non-homologous end-joining(1, 10.00%) |
| Endocrine resistance(1, 2.27%) |
| Rheumatoid arthritis(1, 4.76%) |
| Endometrial cancer(1, 3.85%) |
| Pyrimidine metabolism(1, 3.45%) |
| Acute myeloid leukemia(1, 4.55%) |
| Alanine, aspartate and glutamate metabolism(1, 4.17%) |
| Mucin type O-glycan biosynthesis(1, 25.00%) |
| Bile secretion(1, 4.35%) |
| Quorum sensing(1, 20.00%) |
| Bacterial secretion system(1, 33.33%) |
| Renin-angiotensin system(1, 14.29%) |
| Hematopoietic cell lineage(1, 12.50%) |
| Indole alkaloid biosynthesis(1, 100.00%) |
| Isoquinoline alkaloid biosynthesis(1, 12.50%) |
| Betalain biosynthesis(1, 100.00%) |
| Ascorbate and aldarate metabolism(1, 11.11%) |
| Thiamine metabolism(1, 20.00%) |
| Pentose phosphate pathway(1, 6.25%) |
| Starch and sucrose metabolism(1, 4.76%) |
| Glycosaminoglycan degradation(1, 12.50%) |
| Glycosphingolipid biosynthesis - ganglio series(1, 25.00%) |
| Propanoate metabolism(1, 4.35%) |
| African trypanosomiasis(1, 20.00%) |
| SNARE interactions in vesicular transport(1, 4.76%) |
| Arachidonic acid metabolism(1, 6.25%) |
| alpha-Linolenic acid metabolism(1, 25.00%) |
| Degradation of aromatic compounds(1, 33.33%) |
| Naphthalene degradation(1, 50.00%) |
| Two-component system(1, 7.69%) |
| Limonene and pinene degradation(1, 100.00%) |
| Aldosterone-regulated sodium reabsorption(1, 8.33%) |
| Primary bile acid biosynthesis(1, 25.00%) |
| Streptomycin biosynthesis(1, 25.00%) |
| Neomycin, kanamycin and gentamicin biosynthesis(1, 100.00%) |
| Cyanoamino acid metabolism(1, 25.00%) |

Pathway Sorted by Name

| Pathway  Pathway |
| --- |
| ABC transporters(1, 5.88%) |
| AGE-RAGE signaling pathway in diabetic complications(2, 5.88%) |
| AMPK signaling pathway(3, 5.45%) |
| Acute myeloid leukemia(1, 4.55%) |
| Adherens junction(2, 5.56%) |
| Adipocytokine signaling pathway(3, 12.00%) |
| Adrenergic signaling in cardiomyocytes(3, 6.98%) |
| African trypanosomiasis(1, 20.00%) |
| Alanine, aspartate and glutamate metabolism(1, 4.17%) |
| Alcoholism(5, 10.64%) |
| Aldosterone synthesis and secretion(2, 5.56%) |
| Aldosterone-regulated sodium reabsorption(1, 8.33%) |
| Amino sugar and nucleotide sugar metabolism(5, 16.67%) |
| Aminoacyl-tRNA biosynthesis(2, 7.69%) |
| Amoebiasis(2, 6.67%) |
| Amphetamine addiction(3, 10.71%) |
| Antifolate resistance(1, 7.69%) |
| Antigen processing and presentation(2, 12.50%) |
| Apelin signaling pathway(1, 1.89%) |
| Apoptosis(6, 10.71%) |
| Apoptosis - fly(2, 4.88%) |
| Apoptosis - multiple species(1, 6.25%) |
| Arachidonic acid metabolism(1, 6.25%) |
| Arginine and proline metabolism(2, 8.33%) |
| Arrhythmogenic right ventricular cardiomyopathy(2, 10.00%) |
| Ascorbate and aldarate metabolism(1, 11.11%) |
| Autophagy - animal(3, 3.75%) |
| Axon guidance(1, 1.52%) |
| Axon regeneration(3, 6.25%) |
| B cell receptor signaling pathway(2, 8.70%) |
| Bacterial secretion system(1, 33.33%) |
| Basal cell carcinoma(1, 3.45%) |
| Basal transcription factors(1, 3.23%) |
| Base excision repair(1, 4.17%) |
| Betalain biosynthesis(1, 100.00%) |
| Bile secretion(1, 4.35%) |
| Biosynthesis of amino acids(4, 9.52%) |
| Biosynthesis of cofactors(8, 9.76%) |
| Biosynthesis of unsaturated fatty acids(4, 36.36%) |
| Breast cancer(2, 3.57%) |
| Butanoate metabolism(1, 7.14%) |
| C-type lectin receptor signaling pathway(1, 3.12%) |
| Calcium signaling pathway(5, 7.04%) |
| Carbohydrate digestion and absorption(2, 14.29%) |
| Carbon metabolism(7, 9.59%) |
| Cardiac muscle contraction(2, 7.69%) |
| Cell adhesion molecules(2, 10.00%) |
| Cell cycle(10, 13.70%) |
| Cell cycle - yeast(8, 13.11%) |
| Cellular senescence(3, 4.48%) |
| Central carbon metabolism in cancer(1, 3.33%) |
| Chagas disease(1, 2.86%) |
| Chemical carcinogenesis(4, 26.67%) |
| Chemokine signaling pathway(4, 7.14%) |
| Chloroalkane and chloroalkene degradation(2, 66.67%) |
| Cholesterol metabolism(2, 15.38%) |
| Choline metabolism in cancer(5, 13.51%) |
| Cholinergic synapse(3, 7.14%) |
| Chronic myeloid leukemia(2, 6.25%) |
| Circadian entrainment(1, 2.70%) |
| Cocaine addiction(2, 9.09%) |
| Colorectal cancer(2, 5.26%) |
| Complement and coagulation cascades(1, 7.14%) |
| Cortisol synthesis and secretion(1, 4.17%) |
| Cushing syndrome(4, 5.80%) |
| Cyanoamino acid metabolism(1, 25.00%) |
| Cysteine and methionine metabolism(4, 12.90%) |
| Cytosolic DNA-sensing pathway(2, 9.09%) |
| DNA replication(1, 3.12%) |
| Degradation of aromatic compounds(1, 33.33%) |
| Dilated cardiomyopathy(2, 6.90%) |
| Dopaminergic synapse(3, 6.67%) |
| Dorso-ventral axis formation(2, 9.09%) |
| Drug metabolism - cytochrome P450(4, 44.44%) |
| Drug metabolism - other enzymes(2, 9.09%) |
| ECM-receptor interaction(2, 9.09%) |
| EGFR tyrosine kinase inhibitor resistance(3, 10.34%) |
| Endocrine and other factor-regulated calcium reabsorption(2, 10.00%) |
| Endocrine resistance(1, 2.27%) |
| Endometrial cancer(1, 3.85%) |
| Epithelial cell signaling in Helicobacter pylori infection(1, 3.12%) |
| Epstein-Barr virus infection(4, 5.63%) |
| ErbB signaling pathway(2, 6.06%) |
| Estrogen signaling pathway(5, 10.42%) |
| Ether lipid metabolism(2, 12.50%) |
| Fat digestion and absorption(1, 7.69%) |
| Fatty acid degradation(3, 11.54%) |
| Fatty acid elongation(1, 7.69%) |
| Fatty acid metabolism(4, 12.12%) |
| Fc epsilon RI signaling pathway(2, 9.52%) |
| Fc gamma R-mediated phagocytosis(2, 5.56%) |
| Fluid shear stress and atherosclerosis(3, 6.82%) |
| Focal adhesion(4, 5.13%) |
| Folate biosynthesis(4, 23.53%) |
| FoxO signaling pathway(4, 7.14%) |
| Fructose and mannose metabolism(5, 26.32%) |
| GABAergic synapse(1, 3.23%) |
| Galactose metabolism(1, 5.56%) |
| Gap junction(2, 5.88%) |
| Gastric acid secretion(1, 3.85%) |
| Gastric cancer(3, 5.45%) |
| Glioma(2, 7.41%) |
| Glucagon signaling pathway(2, 4.65%) |
| Glutamatergic synapse(1, 2.38%) |
| Glutathione metabolism(4, 16.67%) |
| Glycerolipid metabolism(3, 11.11%) |
| Glycerophospholipid metabolism(2, 5.00%) |
| Glycine, serine and threonine metabolism(4, 14.29%) |
| Glycolysis / Gluconeogenesis(4, 12.50%) |
| Glycosaminoglycan degradation(1, 12.50%) |
| Glycosphingolipid biosynthesis - ganglio series(1, 25.00%) |
| Glycosphingolipid biosynthesis - globo and isoglobo series(2, 33.33%) |
| Glycosphingolipid biosynthesis - lacto and neolacto series(2, 18.18%) |
| Glyoxylate and dicarboxylate metabolism(3, 12.00%) |
| GnRH secretion(1, 5.26%) |
| GnRH signaling pathway(3, 7.50%) |
| Growth hormone synthesis, secretion and action(4, 8.00%) |
| HIF-1 signaling pathway(2, 4.88%) |
| Hedgehog signaling pathway(1, 4.35%) |
| Hedgehog signaling pathway - fly(1, 4.76%) |
| Hematopoietic cell lineage(1, 12.50%) |
| Hepatitis B(7, 12.07%) |
| Hepatitis C(4, 10.00%) |
| Hepatocellular carcinoma(5, 7.46%) |
| Herpes simplex virus 1 infection(4, 7.27%) |
| Hippo signaling pathway(2, 3.51%) |
| Hippo signaling pathway - fly(2, 4.55%) |
| Histidine metabolism(2, 16.67%) |
| Human T-cell leukemia virus 1 infection(8, 8.89%) |
| Human cytomegalovirus infection(4, 5.26%) |
| Human immunodeficiency virus 1 infection(3, 3.80%) |
| Hypertrophic cardiomyopathy(2, 8.33%) |
| IL-17 signaling pathway(2, 7.41%) |
| Indole alkaloid biosynthesis(1, 100.00%) |
| Inflammatory mediator regulation of TRP channels(1, 3.03%) |
| Influenza A(6, 13.64%) |
| Inositol phosphate metabolism(1, 2.78%) |
| Insect hormone biosynthesis(3, 30.00%) |
| Insulin resistance(3, 6.67%) |
| Insulin secretion(2, 6.06%) |
| Insulin signaling pathway(2, 3.45%) |
| Isoquinoline alkaloid biosynthesis(1, 12.50%) |
| JAK-STAT signaling pathway(2, 9.52%) |
| Kaposi sarcoma-associated herpesvirus infection(2, 3.85%) |
| Legionellosis(3, 13.04%) |
| Leishmaniasis(2, 9.52%) |
| Leukocyte transendothelial migration(1, 3.57%) |
| Limonene and pinene degradation(1, 100.00%) |
| Long-term depression(1, 4.55%) |
| Long-term potentiation(1, 4.00%) |
| Longevity regulating pathway(2, 4.17%) |
| Longevity regulating pathway - worm(2, 5.13%) |
| Lysine degradation(3, 8.33%) |
| Lysosome(6, 9.52%) |
| MAPK signaling pathway(2, 2.22%) |
| MAPK signaling pathway - fly(1, 1.52%) |
| MAPK signaling pathway - yeast(2, 6.90%) |
| Malaria(1, 9.09%) |
| Measles(4, 9.09%) |
| Meiosis - yeast(6, 12.77%) |
| Melanogenesis(3, 7.50%) |
| Metabolism of xenobiotics by cytochrome P450(4, 44.44%) |
| Methane metabolism(3, 18.75%) |
| MicroRNAs in cancer(4, 6.67%) |
| Mineral absorption(2, 12.50%) |
| Mismatch repair(1, 5.00%) |
| Mitophagy - animal(1, 2.70%) |
| Mitophagy - yeast(1, 5.88%) |
| Morphine addiction(1, 3.12%) |
| Mucin type O-glycan biosynthesis(1, 25.00%) |
| N-Glycan biosynthesis(1, 2.86%) |
| NF-kappa B signaling pathway(1, 3.85%) |
| NOD-like receptor signaling pathway(3, 6.00%) |
| Naphthalene degradation(1, 50.00%) |
| Natural killer cell mediated cytotoxicity(2, 9.09%) |
| Necroptosis(3, 8.33%) |
| Neomycin, kanamycin and gentamicin biosynthesis(1, 100.00%) |
| Neuroactive ligand-receptor interaction(1, 1.67%) |
| Neurotrophin signaling pathway(2, 3.92%) |
| Non-alcoholic fatty liver disease(2, 2.38%) |
| Non-homologous end-joining(1, 10.00%) |
| Non-small cell lung cancer(2, 8.00%) |
| Notch signaling pathway(1, 4.00%) |
| Nucleotide excision repair(1, 2.86%) |
| O-Antigen nucleotide sugar biosynthesis(1, 16.67%) |
| Olfactory transduction(1, 5.88%) |
| One carbon pool by folate(5, 45.45%) |
| Oocyte meiosis(5, 8.33%) |
| Osteoclast differentiation(2, 7.14%) |
| Other glycan degradation(2, 20.00%) |
| Other types of O-glycan biosynthesis(2, 13.33%) |
| Oxytocin signaling pathway(2, 3.51%) |
| PD-L1 expression and PD-1 checkpoint pathway in cancer(2, 6.45%) |
| PI3K-Akt signaling pathway(7, 6.80%) |
| PPAR signaling pathway(3, 10.71%) |
| Pancreatic secretion(2, 4.88%) |
| Pantothenate and CoA biosynthesis(1, 9.09%) |
| Parathyroid hormone synthesis, secretion and action(3, 6.67%) |
| Pathogenic Escherichia coli infection(4, 6.25%) |
| Pentose and glucuronate interconversions(1, 7.14%) |
| Pentose phosphate pathway(1, 6.25%) |
| Peroxisome(8, 17.39%) |
| Phagosome(2, 3.92%) |
| Phenylalanine metabolism(2, 20.00%) |
| Phosphatidylinositol signaling system(2, 4.76%) |
| Phospholipase D signaling pathway(2, 3.77%) |
| Phosphonate and phosphinate metabolism(2, 66.67%) |
| Phototransduction - fly(1, 5.88%) |
| Plant-pathogen interaction(2, 18.18%) |
| Platelet activation(1, 2.17%) |
| Platinum drug resistance(4, 14.29%) |
| Porphyrin and chlorophyll metabolism(2, 11.76%) |
| Primary bile acid biosynthesis(1, 25.00%) |
| Progesterone-mediated oocyte maturation(4, 8.51%) |
| Prolactin signaling pathway(2, 7.41%) |
| Propanoate metabolism(1, 4.35%) |
| Prostate cancer(5, 12.82%) |
| Protein digestion and absorption(4, 12.12%) |
| Protein export(5, 23.81%) |
| Proteoglycans in cancer(5, 5.38%) |
| Purine metabolism(3, 4.48%) |
| Pyrimidine metabolism(1, 3.45%) |
| Pyruvate metabolism(1, 4.17%) |
| Quorum sensing(1, 20.00%) |
| RIG-I-like receptor signaling pathway(2, 11.76%) |
| Rap1 signaling pathway(2, 2.70%) |
| Ras signaling pathway(2, 2.60%) |
| Regulation of actin cytoskeleton(3, 4.11%) |
| Relaxin signaling pathway(4, 7.27%) |
| Renal cell carcinoma(1, 3.23%) |
| Renin-angiotensin system(1, 14.29%) |
| Retinol metabolism(2, 18.18%) |
| Retrograde endocannabinoid signaling(2, 2.94%) |
| Rheumatoid arthritis(1, 4.76%) |
| Ribosome biogenesis in eukaryotes(2, 3.12%) |
| SNARE interactions in vesicular transport(1, 4.76%) |
| Salivary secretion(1, 2.78%) |
| Salmonella infection(3, 2.97%) |
| Selenocompound metabolism(2, 22.22%) |
| Serotonergic synapse(2, 5.56%) |
| Shigellosis(2, 1.94%) |
| Signaling pathways regulating pluripotency of stem cells(5, 9.62%) |
| Small cell lung cancer(5, 13.16%) |
| Sphingolipid signaling pathway(2, 4.55%) |
| Spinocerebellar ataxia(3, 3.61%) |
| Starch and sucrose metabolism(1, 4.76%) |
| Streptomycin biosynthesis(1, 25.00%) |
| Systemic lupus erythematosus(2, 12.50%) |
| T cell receptor signaling pathway(2, 5.56%) |
| TGF-beta signaling pathway(1, 3.03%) |
| TNF signaling pathway(3, 8.11%) |
| Th1 and Th2 cell differentiation(2, 8.70%) |
| Th17 cell differentiation(2, 8.00%) |
| Thiamine metabolism(1, 20.00%) |
| Thyroid hormone signaling pathway(2, 3.51%) |
| Thyroid hormone synthesis(5, 15.15%) |
| Tight junction(4, 6.06%) |
| Toll and Imd signaling pathway(1, 3.45%) |
| Toll-like receptor signaling pathway(1, 3.45%) |
| Toxoplasmosis(3, 8.33%) |
| Transcriptional misregulation in cancer(3, 4.17%) |
| Tryptophan metabolism(3, 12.00%) |
| Tuberculosis(3, 6.00%) |
| Two-component system(1, 7.69%) |
| Type II diabetes mellitus(1, 5.56%) |
| Tyrosine metabolism(3, 16.67%) |
| Ubiquitin mediated proteolysis(6, 7.32%) |
| VEGF signaling pathway(1, 4.55%) |
| Valine, leucine and isoleucine degradation(1, 2.86%) |
| Various types of N-glycan biosynthesis(2, 7.69%) |
| Vascular smooth muscle contraction(3, 6.98%) |
| Vasopressin-regulated water reabsorption(2, 8.70%) |
| Vibrio cholerae infection(1, 3.33%) |
| Viral carcinogenesis(9, 11.39%) |
| Vitamin digestion and absorption(1, 12.50%) |
| Wnt signaling pathway(3, 4.17%) |
| Yersinia infection(2, 3.28%) |
| alpha-Linolenic acid metabolism(1, 25.00%) |
| beta-Alanine metabolism(3, 17.65%) |
| cAMP signaling pathway(4, 5.88%) |
| cGMP-PKG signaling pathway(4, 6.78%) |
| mRNA surveillance pathway(1, 1.75%) |
| mTOR signaling pathway(4, 4.60%) |
| p53 signaling pathway(5, 16.67%) |
