## Supplementary material for "The transcriptomic signature of obligate parthenogenesis": Supplementary Material-7-22.docx

**Supplementary Table S1.** List of obligately parthenogenetic (OP) *Daphnia*, cyclically parthenogenetic (CP) *D. pulex*, and CP *D. pulicaria* used in this study.

| **Isolates** | **Location** |
| --- | --- |
| **OP *Daphnia*** |  |
| Maine344-1 | 42°99, -76°01, Appledore Island, Maine |
| MC08 | 46°22, -81°15, McCharles Lake, Sudbury, Ontario |
| **CP *D. pulex*** |  |
| POVI4 | 42°45, -85°21, Battle Creek, Michigan |
| PA42 | 40°13, -87°20, Portland Arch, Indiana |
| **CP *D. pulicaria*** |  |
| AroMoose | 44°50, -69°16, Sebasticook Lake, Maine |
| LK16 | Lab generated inbred line |

**Supplementary Table 2.** Results of Gene Ontology (GO) term enrichment analysis for underdominant genes.

| **GO ID** | **GO Term** | **Identified genes** | **Expected** | **p.adjust** |
| --- | --- | --- | --- | --- |
| GO:0007346 | regulation of mitotic cell cycle | 5 | 0.21 | 0.003537 |
| GO:0007049 | cell cycle | 9 | 1.23 | 0.003537 |
| GO:0010564 | regulation of cell cycle process | 5 | 0.24 | 0.003537 |
| GO:1903047 | mitotic cell cycle process | 6 | 0.47 | 0.004421 |
| GO:0000278 | mitotic cell cycle | 6 | 0.5 | 0.004421 |
| GO:0044770 | cell cycle phase transition | 4 | 0.15 | 0.004421 |
| GO:0044772 | mitotic cell cycle phase transition | 4 | 0.15 | 0.004421 |
| GO:1901987 | regulation of cell cycle phase transition | 4 | 0.15 | 0.004421 |
| GO:1901990 | regulation of mitotic cell cycle phase transition | 4 | 0.15 | 0.004421 |
| GO:0007091 | metaphase/anaphase transition of mitotic cell cycle | 3 | 0.09 | 0.011274 |
| GO:0010965 | regulation of mitotic sister chromatid separation | 3 | 0.09 | 0.011274 |
| GO:0030071 | regulation of mitotic metaphase/anaphase transition | 3 | 0.09 | 0.011274 |
| GO:0033045 | regulation of sister chromatid segregation | 3 | 0.09 | 0.011274 |
| GO:0033047 | regulation of mitotic sister chromatid segregation | 3 | 0.09 | 0.011274 |
| GO:0044784 | metaphase/anaphase transition of cell cycle | 3 | 0.09 | 0.011274 |
| GO:0051304 | chromosome separation | 3 | 0.09 | 0.011274 |
| GO:0051306 | mitotic sister chromatid separation | 3 | 0.09 | 0.011274 |
| GO:0051983 | regulation of chromosome segregation | 3 | 0.09 | 0.011274 |
| GO:1902099 | regulation of metaphase/anaphase transition | 3 | 0.09 | 0.011274 |
| GO:1905818 | regulation of chromosome separation | 3 | 0.09 | 0.011274 |
| GO:0033043 | regulation of organelle organization | 5 | 0.47 | 0.014417 |
| GO:0007088 | regulation of mitotic nuclear division | 3 | 0.1 | 0.014417 |
| GO:0051783 | regulation of nuclear division | 3 | 0.1 | 0.014417 |
| GO:0070507 | regulation of microtubule cytoskeleton organization | 2 | 0.02 | 0.01658 |
| GO:0022402 | cell cycle process | 6 | 0.8 | 0.01857 |
| GO:0000070 | mitotic sister chromatid segregation | 4 | 0.29 | 0.020133 |
| GO:0000819 | sister chromatid segregation | 4 | 0.29 | 0.020133 |
| GO:0098813 | nuclear chromosome segregation | 4 | 0.29 | 0.020133 |
| GO:0006270 | DNA replication initiation | 3 | 0.12 | 0.022869 |
| GO:0051726 | regulation of cell cycle | 5 | 0.57 | 0.026528 |
| GO:0033044 | regulation of chromosome organization | 3 | 0.13 | 0.027812 |
| GO:0007059 | chromosome segregation | 4 | 0.33 | 0.030051 |
| GO:0032886 | regulation of microtubule-based process | 2 | 0.03 | 0.03517 |
| GO:0051128 | regulation of cellular component organization | 5 | 0.62 | 0.036002 |
| GO:0140014 | mitotic nuclear division | 4 | 0.35 | 0.036002 |
| GO:0000280 | nuclear division | 4 | 0.38 | 0.046424 |
| GO:0007094 | mitotic spindle assembly checkpoint | 2 | 0.04 | 0.046424 |
| GO:0031577 | spindle checkpoint | 2 | 0.04 | 0.046424 |
| GO:0033046 | negative regulation of sister chromatid segregation | 2 | 0.04 | 0.046424 |
| GO:0033048 | negative regulation of mitotic sister chromatid segregation | 2 | 0.04 | 0.046424 |
| GO:0045839 | negative regulation of mitotic nuclear division | 2 | 0.04 | 0.046424 |
| GO:0045841 | negative regulation of mitotic metaphase/anaphase transition | 2 | 0.04 | 0.046424 |
| GO:0051784 | negative regulation of nuclear division | 2 | 0.04 | 0.046424 |
| GO:0051985 | negative regulation of chromosome segregation | 2 | 0.04 | 0.046424 |
| GO:0071173 | spindle assembly checkpoint | 2 | 0.04 | 0.046424 |
| GO:0071174 | mitotic spindle checkpoint | 2 | 0.04 | 0.046424 |
| GO:1902100 | negative regulation of metaphase/anaphase transition | 2 | 0.04 | 0.046424 |
| GO:1905819 | negative regulation of chromosome separation | 2 | 0.04 | 0.046424 |
| GO:2000816 | negative regulation of mitotic sister chromatid separation | 2 | 0.04 | 0.046424 |
| GO:2001251 | negative regulation of chromosome organization | 2 | 0.04 | 0.046424 |

**Supplementary Table 3.** Results of Gene Ontology (GO) term enrichment analysis for overdominant genes.

| **GO ID** | **GO Term** | **Identified genes** | **Expected** | **p.adjust** |
| --- | --- | --- | --- | --- |
| GO:0006486 | protein glycosylation | 13 | 2.77 | 0.003758 |
| GO:0043413 | macromolecule glycosylation | 13 | 2.77 | 0.003758 |
| GO:0070085 | glycosylation | 13 | 2.77 | 0.003758 |
| GO:0009101 | glycoprotein biosynthetic process | 13 | 2.93 | 0.004908 |
| GO:0009100 | glycoprotein metabolic process | 13 | 2.97 | 0.004908 |
| GO:1901564 | organonitrogen compound metabolic process | 73 | 48 | 0.005085 |
| GO:0055114 | oxidation-reduction process | 27 | 11.18 | 0.006158 |
| GO:1901135 | carbohydrate derivative metabolic process | 24 | 9.52 | 0.00829 |
| GO:1901137 | carbohydrate derivative biosynthetic process | 16 | 5.04 | 0.014369 |
| GO:0006635 | fatty acid beta-oxidation | 3 | 0.1 | 0.025533 |
| GO:0019395 | fatty acid oxidation | 3 | 0.12 | 0.04145 |
| GO:0034440 | lipid oxidation | 3 | 0.12 | 0.04145 |

**Supplementary File 1.** The list of genes that are underdominant in all obligately parthenogenetic *Daphnia* genotypes.

**Supplementary File 2.** The list of genes that are overdominant in all obligately parthenogenetic *Daphnia* genotypes.

**Supplementary File 3.** The list of genes that are underdominant in all obligately parthenogenetic *Daphnia* genotypes and do not have expression changes at the within- and between-species levels.

**Supplementary File 4.** The list of genes that are overdominant in all obligately parthenogenetic *Daphnia* genotypes and do not have expression changes at the within- and between-species levels.

**Supplementary File 5.** The original list of underdominant (red color) and overdominant (green) genes.

**Supplementary File 6.** The select list of underdominant (red color) and overdominant (green) genes that do not have significant expression changes at within- and between-species levels.
